# Genetic and microbial diversity of the invasive mosquito vector species *Culex tritaeniorhynchus* across its extensive inter-continental geographic range

**DOI:** 10.1101/2022.02.10.479990

**Authors:** Claire L. Jeffries, Luciano M. Tantely, Perparim Kadriaj, Marcus S. C. Blagrove, Ioanna Lytra, James Orsborne, Hasan M. Al-Amin, Abdul Rahim Mohammed, Mohammad Shafiul Alam, Romain Girod, Yaw A. Afrane, Silvia Bino, Vincent Robert, Sebastien Boyer, Matthew Baylis, Enkelejda Velo, Grant L. Hughes, Thomas Walker

## Abstract

*Culex (Cx.) tritaeniorhynchus* is a mosquito species with an extensive and expanding inter-continental geographic distribution, currently reported in over 50 countries, across Asia, Africa, the Middle East, Europe and now Australia. It is an important vector of medical and veterinary concern, capable of transmitting multiple arboviruses which cause significant morbidity and mortality in human and animal populations. In regions endemic for Japanese encephalitis virus (JEV) in Asia, *Cx. tritaeniorhynchus* is considered the major vector and this species has also been shown to contribute to the transmission of several other significant zoonotic arboviruses, including Rift Valley fever virus and West Nile virus.

Significant variation in vectorial capacity can occur between different vector populations. Obtaining knowledge of a species from across its geographic range is crucial to understanding its significance for pathogen transmission across diverse environments and localities. Vectorial capacity can be influenced by factors including the mosquito genetic background, composition of the microbiota associated with the mosquito and the co-infection of human or animal pathogens. In addition to enhancing information on vector surveillance and potential risks for pathogen transmission, determining the genetic and microbial diversity of distinct populations of a vector species is also critical for the development and application of effective control strategies.

In this study, multiple geographically dispersed populations of *Cx. tritaeniorhynchus* from countries within Europe, Africa, Eurasia and Asia were sampled. Molecular analysis demonstrated a high level of genetic and microbial diversity within and between populations, including genetic divergence in the mosquito *CO1* gene, as well as diverse microbiomes identified by *16S rRNA* gene amplicon sequencing. Evidence for the detection of the endosymbiotic bacteria *Wolbachia* in some populations was confirmed using *Wolbachia*-specific PCR detection and sequencing of *Wolbachia* MLST genes; in addition to PCR-based detection of insect-specific viruses. Laboratory vector competence showed *Cx. tritaeniorhynchus* from a Greek population are likely to be competent vectors of JEV. This study expands understanding of the diversity of *Cx. tritaeniorhynchus* across its inter-continental range, highlights the need for a greater focus on this invasive vector species and helps to inform potential future directions for development of vector control strategies.

## 1 Introduction

The invasive mosquito vector species *Culex (Cx.) tritaeniorhynchus* (Giles 1901) has a wide and expansive distribution which includes populations in over 50 countries. Ranging across Asia, the Middle East and Africa (1), it has in recent decades been additionally reported in Europe (2), Eurasia (3), Cape Verde off western Africa (4) and in 2020 it was recorded for the first time in Australia (5). It is a vector of significant medical and veterinary importance; the major vector of Japanese encephalitis virus (JEV) (1), and capable of transmitting several other significant zoonotic arboviruses, including Rift Valley fever virus (RVFV) (6) and West Nile virus (WNV) (7–10), with mosquito and viral geographic distributions extensively overlapping.

JEV (Family: *Flaviviridae*, Genus: *Flavivirus*) is transmitted to humans, birds, pigs, and other vertebrates through infectious mosquito blood-feeding (11). Human disease ranges from asymptomatic or mild flu-like illness, to severe encephalitic disease (JE) and death. Case-fatality rates from JE are 20-30%, with 30-50% of survivors suffering serious, often long-lasting, neurological sequelae (12). Furthermore, JEV causes reproductive problems and abortion in pigs and neurological disease in horses (11). As the major vector in most areas of Asia and the Pacific where JEV is endemic, *Cx. tritaeniorhynchus* is a highly important contributor to viral transmission which leads to an estimated 50,000 – 175,000 human JE disease cases annually (13,14). Estimates suggest JE presentations account for 1% of total viral infections, indicating overall occurrence of human JEV infections could be in the region of 5 – 17.5 million each year, with almost 4 billion people living in 24 endemic countries at risk (1,11). There have also been recent cases of JEV detection in mosquitoes and birds in Italy, Europe (15,16) and an autochthonous human case in Angola, Africa (17), highlighting the possibilities of future viral spread and establishment in novel regions and naïve populations (11,18,19).

RVFV (Family: *Phenuiviridae*, Genus: *Phlebovirus*) is distributed across Africa, and now the Middle East, in 32 endemic countries. *Cx. tritaeniorhynchus* was a major vector of RVFV during an epidemic in Saudi Arabia in 2000 (6); the first occurrence of RVFV outside of Africa, leading to nearly 900 human cases of infection and 124 deaths, with further infections in neighbouring Yemen (20,21). Approximately 780 million people live in endemic countries and are potentially at-risk from RVFV, with high variability in annual incidence due to explosive outbreaks and epidemics (12).

Human disease ranges from asymptomatic or mild flu-like illness, to severe disease resulting in hepatitis, encephalitis, retinitis or haemorrhagic fever. Case-fatality rates can be 10-20%, rising up to 50% in haemorrhagic manifestations, and survivors can have long-lasting health consequences (21). Veterinary disease presentations include reproductive problems, abortion and death in ruminants, with “abortion storms” often a characteristic of outbreaks. In addition to the suffering of animals, the economic and food security risks from livestock losses can be significant (12). There is growing concern for future increased occurrence, re-emergence or expansions of RVFV in several regions, with serious potential consequences for human and animal health (11,22,23).

WNV (Family: *Flaviviridae*, Genus: *Flavivirus*) is globally distributed and endemic to all continents except Antarctica; putting populations worldwide at risk of infection (24). Human disease occurs as a spectrum from asymptomatic or mild flu-like illness to severe neurological syndromes and death (11). Estimates of global annual incidence are unknown due to asymptomatic infections, variable detection and reporting, apparent variation in virulence of WNV lineages, and wide fluctuations in outbreak occurrence year-on-year (25). The human case-fatality rate, however, is approximately 10% of neurological disease cases, with survivors suffering long-term health consequences and morbidities. Veterinary disease manifestations include neurological syndromes and death in some avian species and horses (26). *Cx. tritaeniorhynchus* has been implicated as a competent vector of WNV in certain countries (7,8,10,27), however, its capacity and contribution to transmission appears to be under-studied, particularly considering that the small amount of vector competence data which is available indicates high susceptibility to WNV infection, including when tested comparatively in Pakistan (7); exhibiting an even a greater susceptibility to infection than *Cx. quinquefasciatus* – generally considered one of the major global WNV vectors (28).

In addition to these major arboviruses, *Cx. tritaeniorhynchus* has also been implicated (through laboratory experiments or field studies), as a competent or potential vector of: Sindbis virus (29) and Getah virus (10,30) (Family: *Togaviridae*, Genus: *Alphavirus*); Bagaza virus (31,32) and Tembusu virus (33) (Genus: *Flavivirus*); Batai (Chittoor) virus (34), Manzanilla virus (35), Cat Que virus (36), and Akabane virus (37) (Family: *Peribunyaviridae*, Genus: *Orthobunyavirus*); Banna virus (38) (Family: *Reoviridae*, Genus: *Seadornavirus*); Mengovirus (Cardiovirus A) (8) (Family: *Picornaviridae*, Genus: *Cardiovirus*); Chandipura virus (39) (Family: *Rhabdoviridae*, Genus: *Vesiculovirus*); and the filarial nematode parasite *Dirofilaria immitis* (40).

Inter-population variability in the vectorial capacity of a vector species can be influenced by multiple endogenous features, including mosquito genetic diversity, any host blood-feeding preference, the composition of their natural symbiotic microbiota, and the presence of other infecting microbes (41–46). The complexity and interaction of vectorial capacity determinants and transmission networks emphasizes the value of obtaining information on genetic and microbial variation from diverse populations (47,48). This not only provides vital information on intrinsic components influencing transmission potential in different locations, but with development of genetic and microbial biocontrol approaches, can also highlight potential transmission-reduction strategies; through mosquito population reduction, vector refractoriness or direct pathogen interference (47,49). Laboratory vector competence studies can measure the capability and efficiency of a vector population, under experimental conditions, to acquire (becoming infected after a feed) and go on to transmit a pathogen (producing infectious saliva during subsequent feeding). Although it is unfortunately too simplistic to directly relate experimental results to the transmission risk from, or vectorial capacity of, mosquitoes in the wild; vector competence measurements form a component part of the wider inter-connected elements of vectorial capacity. Such experimental assessments provide important information on the potential for onward transmission, can indicate the functional effects of intrinsic characteristics and help to elucidate the potential effects of any variation observed (50).

Beside pathogenic arboviruses, there are numerous viruses which are associated with invertebrate vector species, but which appear to be incapable of replicating in vertebrate cells (invertebrate-specific), or where no pathogenicity has so far been detected in vertebrate hosts (51). Several invertebrate-associated viruses are closely related to pathogenic arboviruses (such as those within the *Flavivirus* genus) and are widespread in certain vector populations (41). Their presence in medically important vector species and the potential for co-infection to influence infection dynamics are important considerations for pathogen detection, vectorial capacity and disease control (41,52). Mosquitoes also have associations with a wide diversity of other microbes, some of which appear to have a range of potential effects, roles and functions within their invertebrate hosts (43). Examples include bacteria such as *Wolbachia, Asaia*, *Serratia* and *Pseudomonas* (41,53). Some of these microbes can affect vectorial capacity, either indirectly through influences on mosquito fitness and immunity, or directly through pathogen interference during co-infection (41,43,54).

Although several previous studies have investigated *Cx. tritaeniorhynchus* genetic variation, and some have examined variation in vector competence, they have limitations; pre-dating modern molecular characterization techniques, and/or being confined to populations from within the same region or country, mainly within Asia (10,55–61). In addition to genetic diversity and vector competence, several arthropod-associated viruses have been recovered from *Cx. tritaeniorhynchus* (62–66), and some studies on the native microbiome or associated bacteria have been carried out (67–69), but microbiome composition and the presence of other infecting microbes has not been extensively characterized or compared, within and between populations of *Cx. tritaeniorhynchus*. Expanding the data available for this species is essential to better understand the variation in intrinsic influences on vectorial capacity across diverse populations.

In this study, we obtained geographically dispersed collections from multiple populations spanning four continents. A variety of molecular analyses examined genetic and microbial diversity of this invasive and medically important vector species. Our analysis provided evidence for the presence of novel strains of the endosymbiotic bacteria *Wolbachia*, in addition to the presence of mosquito-only viruses. We also undertook a study of laboratory vector competence for JEV, demonstrating *Cx. tritaeniorhynchus* from European populations have the potential to be competent vectors of JEV. This study provides important data to expand knowledge on the potential role of *Cx. tritaeniorhynchus* in transmission of significant diseases and the possibilities for control strategies.

## 2 Materials and Methods

### 2.1 Mosquito Collections

*Cx. tritaeniorhynchus* specimens were obtained from field-collections in Albania and Greece in Europe, Georgia in Eurasia, Ghana and Madagascar in Africa, and Bangladesh and India in Asia. The geographic distribution of field-collected specimens is shown in Figure 1. The locations, year, GPS co-ordinates, methods of collection and number of specimens collected are shown in Table 1.

**Fig 1.**
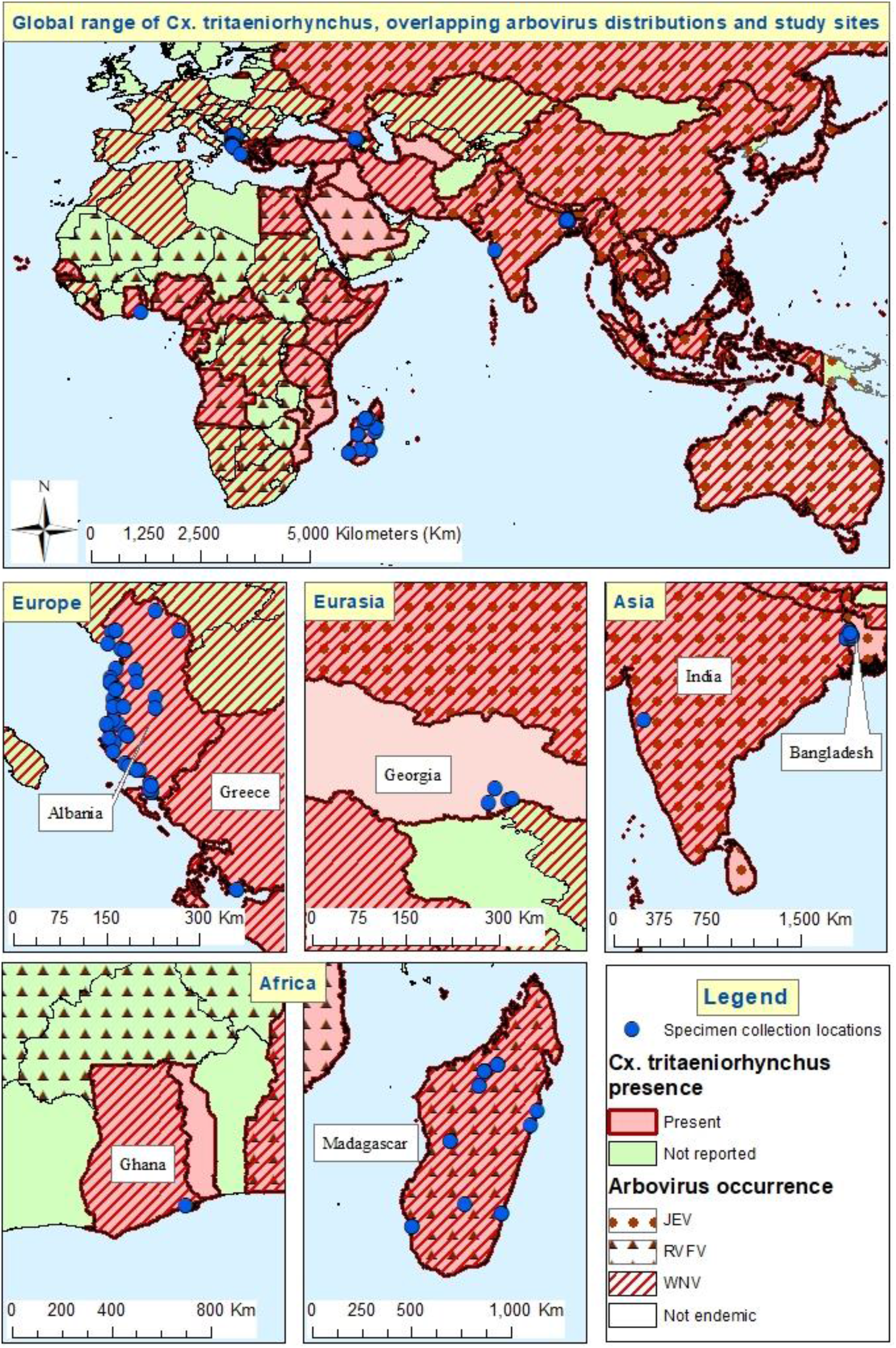
Overlapping viral and mosquito geographic distributions and sampling locations.

**Table 1.**
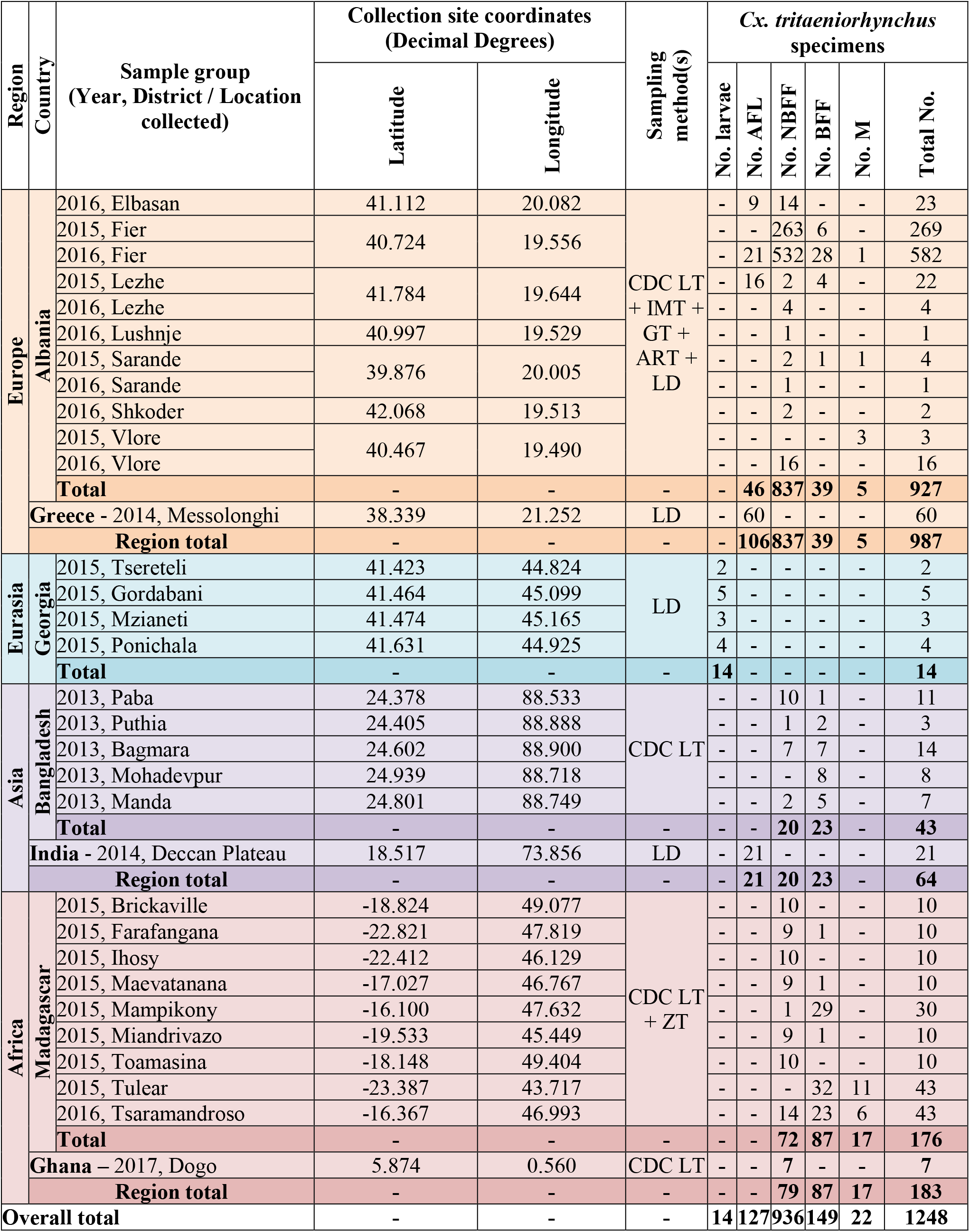
*Cx. tritaeniorhynchus* specimens and collection locations.

#### Collections from Europe and Eurasia

Specimens were obtained from Albania during country-wide entomological surveys in 2015 and 2016, in addition to focused field-work collections of adults and sampling through larval dipping in locations where high densities of *Cx. tritaeniorhynchus* were previously found, particularly in the rural village of Sop in the Fier district in south western Albania. Specimens were preserved in RNAlater, combined with cold temperature storage. In Greece, larval dipping was used to collect live *Cx. tritaeniorhynchus* larvae in 2014 from the irrigated rice fields where this species was previously identified within the Messolonghi district, western Greece (70). Live larvae were then shipped to LSHTM for initiation of a laboratory colony, with specimens from each generation stored for preservation of RNA. Field-collected fourth instar *Cx. tritaeniorhynchus* larvae were obtained from four sites in south eastern Georgia in September 2015; using larval dipping in semi-permanent water bodies with vegetation, then stored in 70% ethanol.

#### Collections from Africa

In Ghana, specimens were collected as adults from the village of Dogo in the Greater Accra region of Ghana in June 2017 as detailed in Orsborne *et al*. 2019 (71). Adult specimens were preserved in RNAlater with cold storage. Sampling in Madagascar was carried out in 2015/2016, from locations spanning the various bioclimatic ecotype zones across Madagascar as described in Jeffries *et al*. 2018 (72). These specimens were also preserved in RNAlater with cold storage to prevent RNA degradation.

#### Collections from Asia

Adult *Cx. tritaeniorhynchus* collections in Bangladesh were carried out in Sept-Nov 2013 from five sites within two districts in the Rajshahi Division in western Bangladesh. Within the district of Rajshahi, mosquitoes were collected from the upazilas (sub-districts) of Paba, Puthia and Bagmara, and within the Naogaon district, specimens were obtained from the upazilas of Manda and Mohadevpur. Samples were then stored dry with desiccant. In India, wild *Cx. tritaeniorhynchus* mosquito eggs and larvae were collected from rice paddy fields in different parts of the Deccan Plateau and used to initiate a laboratory colony with specimens from subsequent generations stored in RNAlater with cold storage.

### 2.2 Morphological Identification, Nucleic Acid Extraction and Molecular Confirmation

Specimens were morphologically identified using keys appropriate for the geographic region from which they were collected (2,70,73–75) and examined for relevant *Cx. tritaeniorhynchus*-specific morphological characteristics, such as the clear white band on the proboscis, entirely dark wings and ringed tarsi (the distal segments of the mosquito legs) (Supplementary Figure S1a) (2). Adult female mosquito physiological status was recorded and if wild-caught blood-fed, then the stage of digestion and time since blood-feeding was approximated using the Sella score method (Supplementary Figure S1b) (76,77).

Following morphological examination, and dependent upon the collection techniques, preservation methods and possibilities for downstream analysis, the relevant nucleic acid extraction methods were employed. All specimens were homogenized using a Qiagen Tissue Lyser II and 3mm stainless steel beads. DNA was extracted from Georgia and Bangladesh specimens using a Qiagen DNeasy Blood and Tissue kit according to the manufacturer’s instructions. Where investigation of viruses was possible from the preservation and physiological status of the specimens, RNA extraction alone, or a modified method for simultaneous RNA and DNA co-extraction utilizing Trizol reagent extraction protocol, prior to column-based extraction of the relevant phase using Qiagen DNeasy or RNeasy kits was used. RNA eluates were converted to cDNA using Reverse Transcription kits according to manufacturer’s instructions (Further details provided in Supplementary Material). To confirm morphological species identification, gDNA or cDNA from a sub-set of specimens was used in broad-specificity barcoding PCRs, followed by Sanger sequencing and phylogenetic analysis (as detailed for genetic diversity analysis below) to confirm the species identification.

### 2.3 Intra- and inter-population genetic diversity

#### 2.3.1 Molecular assays and sequencing strategy

Genetic diversity of *Cx. tritaeniorhynchus* populations was assessed through Sanger sequencing of amplified PCR products from several assays, targeting the mitochondrial gene cytochrome oxidase subunit 1 (*CO1*) (78–81) or the nuclear internal transcribed spacer 2 (ITS2) region (82–84). A large number of *CO1* fragment primer sets have been designed and used for species barcoding and to investigate genetic variation in mosquito species, targeting various regions of the mt *CO1* gene and having been used in different geographic regions. Therefore, preliminary sub-sample testing was carried out to assess; (i) the success of amplification and sequencing for *Cx. tritaeniorhynchus* specimens – trying to minimize the risk of amplification bias due to mutations in primer binding regions, and (ii) the number of comparative sequences with fragment coverage already publicly available for reference – thereby maximizing the level of discrimination and breadth of diversity analysis possible. The primer set designed by Kumar et al. (78) (MTFN 5’- GGATTTGGAAATTGATTAGTTCCTT-3’ and MTRN 5’- AAAAATTTTAATTCCAGTTGGAACAGC-3’) producing a product ~700 base pair (bp) in length was selected for screening a larger number of samples across all populations. A primer combination to amplify the full length of the *CO1* gene (TY-J-1460 5’- TACAATTTATCGCCTAAACTTCAGCC-3’ and TL2-N-3014 (later described as UEA10) 5’- TCCAATGCACTAATCTGCCATATTA-3’), binding at the 5’ and 3’ tRNA respectively to produce a ~1150bp sequence (79,85,86), was also used on selected samples to generate longer sequences. (Supplementary Figure S2 and further details provided in Supplementary Material.)

#### 2.3.2 Consensus sequence and alignment assembly

Sequencing analysis was carried out in MEGA11 (87) as follows. Both chromatograms (forward and reverse traces) from each sample were manually checked, edited, and trimmed as required, followed by alignment by ClustalW and checking to produce consensus sequences. Consensus sequences were used to perform nucleotide BLAST (NCBI) database queries, which informed the building of alignments for each examined target fragment, comprising all consensus sequences generated, alongside relevant reference sequences obtained from GenBank. All mitochondrial *CO1* nucleotide sequences for *Cx. tritaeniorhynchus* available on GenBank (NCBI: txid7178, 992 sequences) were downloaded and aligned with the *CO1* sequences (69 for *Cx. tritaeniorhynchus*) generated in this study. This initial alignment was checked and then edited according to three criteria. Three separate CO1 alignments were constructed to include; (a) all *Cx. tritaeniorhynchus* sequences with coverage of the fragment generated by the Kumar *et al*. primer set (78) (253 sequences, 686 positions) and including concomitant species obtained during field-collections, (b) all *Cx. tritaeniorhynchus CO1* gene full length sequences, maximizing the length (20 sequences, 1538 positions), and (c) comprising all *Cx. tritaeniorhynchus CO1* sequences currently available with sufficient fragment overlap, to balance the length of the alignment but maximize the number of reference sequences included (1007 sequences, 414 positions). Sequences with missing data or nucleotide ambiguities were excluded. The positions of the alignments and primer binding regions according to the *Cx. tritaeniorhynchus* complete mitochondrial genome reference sequence NC_028616 is provided in Supplementary Figure S2.

#### 2.3.3 Phylogenetic tree construction and analysis

Each alignment was examined using the “Find-Best-Fit Maximum Likelihood substitution model” to identify the best options for phylogenetic analysis and tree construction. The model with the lowest Bayesian information criterion (BIC) score from this analysis is considered to describe the substitution pattern the best. Options to model non-uniformity of evolutionary rates among sites using a discrete Gamma distribution (+*G*) with five rate categories and by assuming that a certain fraction of sites is evolutionary invariable (+*I*) were also evaluated during this analysis to highlight the most appropriate model and options to use for construction of each phylogenetic tree. The evolutionary history was then inferred by using the ML method with the most appropriate model and options for each respective tree selected, with details of the methods and parameters used for each specific tree included in the figure legends. The models used in the analysis were the General Time Reversible model (88) (GTR) or the Tamura three-parameter model (89) (T92). The tree with the highest log likelihood is shown. The percentage of trees in which the associated taxa clustered together is shown next to the branches. Initial tree(s) for the heuristic search were obtained automatically by applying Neighbor-Joining and BioNJ algorithms to a matrix of pairwise distances estimated using the Maximum Composite Likelihood (MCL) approach, and then selecting the topology with superior log likelihood value. The trees are drawn to scale, with branch lengths measured in the number of substitutions per site. Codon positions included were 1st+2nd+3rd+Noncoding. All positions containing gaps and missing data were eliminated. The phylogeny test was by Bootstrap method with 1000 replications. Evolutionary analyses were conducted in MEGA11 (87).

#### 2.3.4 Genetic diversity and haplotype analyses

Genetic diversity of the *Cx. tritaeniorhynchus* populations was further assessed through the calculation of genetic diversity metrics, analysis of haplotypes, with the generation of haplotype networks, and pairwise comparison of genetic differentiation, including both study-generated and available reference *CO1* sequences, at the individual, population and regional levels. The *Cx. tritaeniorhynchus CO1* alignments were analyzed using DnaSP V6.12.03 (90) to assess sequence polymorphisms and determine nucleotide and haplotype diversity. Haplotype networks were constructed within PopART (91) using the TCS inference method (92). Intra- and inter-group variation was assessed at the individual, country and regional population levels using Arlequin V3.5.2.2 (93), with analysis of molecular variance (AMOVA) (94) and visualization of outputs in R V3.5.0 (95).

### 2.4 European population colonisation and JEV vector competence experiments

Field-collected *Cx tritaeniorhynchus* larvae (~500) from Messolonghi, Greece were transported to LSHTM for initiation of a colony from wild larvae. A range of techniques were employed to optimize conditions in the insectaries for all life stages of mosquito (detailed in supplementary figure 3). Alongside the Greek *Cx. tritaeniorhynchus* colonisation, multiple shipments of *Cx. tritaeniorhynchus* eggs and larvae from other countries were used to attempt colony initiation from additional source populations for comparative purposes. However, difficulties in establishing and maintaining these colonies prevented comparative experimental data from being obtained. JEV vector competence was assessed on the fourth generation of the Greek *Cx. tritaeniorhynchus* colony at the Liverpool School of Tropical Medicine. Blood meals (heparinized human blood, NHS transfusion service, Speke) containing JEV (strain CNS138-11), to a final concentration of 6 log10 plaque-forming units/mL, were provided for 3 hours, using a Hemotek membrane feeding system and an odorized feeding membrane, to 5-7-day-old adult females from which sugar sources had been withheld for 24 hours. Blood-fed females were incubated at 27°C, 70% humidity, for 14 days prior to collection of saliva using a forced salivation technique (96). The head/thorax and abdomen were separated for each of the 28 surviving females after the 14-day incubation and the dissected body parts were stored for RNA preservation. RNA was extracted from all saliva and body-part samples and tested by JEV-specific real-time PCR analysis (97) to determine infection rates.

### 2.5 Molecular screening for arboviruses and insect-only viruses

A range of molecular methods for arbovirus detection were used, with the samples screened for arboviruses and invertebrate-associated viruses using a combination of broad pan-virus assays (such as Pan-Flavivirus (98), Pan-Alphavirus (99) and Pan-Orthobunyavirus (100) PCRs) and virus specific PCRs, including assays for detection of WNV (101), JEV (97) and RVFV (102). Sequencing was attempted for virus positive PCR products to confirm virus detection and provide sequencing data for phylogenetic analysis.

### 2.6 Microbial diversity through microbiome analysis

#### 2.6.1 *16S rRNA* gene amplicon sequencing

The microbiomes of selected individual mosquitoes were analyzed using barcoded high-throughput amplicon sequencing of the bacterial *16S rRNA* gene. To enable analysis of the differences in microbiome between species (*Cx. tritaeniorhynchus* and concomitant species), physiological status (blood-fed or non-blood-fed) and geographic location (both intra- and inter-country) samples were selected from specific groups for comparison (e.g. gDNA or cDNA, *Cx. tritaeniorhynchus* or concomitant species, extracts from whole body or abdomen, blood-fed or non-blood-fed, country and/or location of collection) (Table 3). Mosquito specimens were surface sterilized prior to extraction, and negative controls comprising both DNA extraction and RNA extraction–Reverse Transcription blanks were also included alongside the samples throughout processing. Sequencing of each extract was achieved using universal *16S rRNA* V3-V4 region primers (FOR: 5’-CCTACGGGNGGCWGCAG-3’, REV: 5’-GGACTACHVGGGTATCTAATCC-3’) (103) in accordance with standard Illumina *16S rRNA* metagenomic sequencing library protocols with the Nextera XT Index Kit v2 used to barcode samples for multiplexing. Sequencing was performed on an Illumina MiSeq, with the MiSeq v2 (500 cycle) reagent kit, with libraries sequenced as 250bp paired-end reads (PE).

**Table 2.** *Cx. tritaeniorhynchus* population genetic diversity metrics.

| Region | Country | n | S | h | Hd | K | Pi | PiJC |
| --- | --- | --- | --- | --- | --- | --- | --- | --- |
| **Asia** | **All (n=11)** | **909** | **137** | **412** | **0.97487** | **9.12217** | **0.02203** | **0.02265** |
|  | **Bangladesh** | 10 | 13 | 9 | 0.97778 | 2.77778 | 0.00671 | 0.00675 |
|  | China | 608 | 116 | 290 | 0.96684 | 10.07545 | 0.02434 | 0.02509 |
|  | **India** | 159 | 77 | 83 | 0.91673 | 3.77351 | 0.00911 | 0.00927 |
|  | Japan | 33 | 40 | 30 | 0.99432 | 12.97159 | 0.03133 | 0.03226 |
|  | Pakistan | 80 | 30 | 31 | 0.84778 | 2.44399 | 0.00590 | 0.00594 |
|  | Singapore | 6 | 9 | 6 | 1.00000 | 3.20000 | 0.00773 | 0.00778 |
|  | South Korea | 3 | 22 | 3 | 1.00000 | 14.66667 | 0.03543 | 0.03653 |
|  | Sri Lanka | 7 | 7 | 5 | 0.85714 | 2.00000 | 0.00483 | 0.00486 |
|  | Vietnam | 1 | N/A |  |  |  |  |  |
|  | Thailand | 1 | N/A |  |  |  |  |  |
|  | Timor Leste | 1 | N/A |  |  |  |  |  |
| **Australia** | **Australia** | **19** | **6** | **4** | **0.73099** | **2.33918** | **0.00565** | **0.00569** |
| **Africa** | **All (n=2)** | **34** | **22** | **19** | **0.95544** | **3.39037** | **0.00819** | **0.00825** |
|  | **Madagascar** | 28 | 16 | 14 | 0.93651 | 2.86508 | 0.00692 | 0.00697 |
|  | **Ghana** | 6 | 6 | 5 | 0.93333 | 2.20000 | 0.00531 | 0.00534 |
| **Middle East** | **All (n=3)** | **4** | **8** | **4** | **1.00000** | **4.50000** | **0.01087** | **0.01098** |
|  | Kuwait | 2 | 1 | 2 | 1.00000 | 1.00000 | 0.00242 | 0.00242 |
|  | Saudi Arabia | 1 | N/A |  |  |  |  |  |
|  | UAE | 1 | N/A |  |  |  |  |  |
| **Eurasia** | **All (n=2)** | **22** | **13** | **8** | **0.85714** | **3.28571** | **0.00794** | **0.00800** |
|  | Turkey | 13 | 12 | 6 | 0.82051 | 3.35897 | 0.00811 | 0.00818 |
|  | **Georgia** | 9 | 2 | 3 | 0.55556 | 0.88889 | 0.00215 | 0.00215 |
| **Europe** | **All (n=2)** | **19** | **3** | **4** | **0.64327** | **0.76023** | **0.00184** | **0.00184** |
|  | **Albania** | 11 | 3 | 4 | 0.60000 | 0.69091 | 0.00167 | 0.00167 |
|  | **Greece** | 8 | 1 | 2 | 0.53571 | 0.53571 | 0.00129 | 0.0013 |
| **ALL SEQUENCES TOTALS** |  | **1007** | **139** | **444** | **0.97864** | **9.16425** | **0.02214** | N/A |

Total number of sites: 414 n: Number of samples; S: Number of variable sites; h: Number of haplotypes; Hd: Haplotype diversity; K: Average number of nucleotide differences; Pi: Nucleotide diversity (per site); PiJC: Nucleotide diversity (Jukes-Cantor)

**Table 3.** Sampling groups and associated information for *16S* microbiome analysis.

Abbreviations: NBFF = Non-blood-fed-female, BFF = Blood-fed-female, gDNA = genomic DNA, cDNA = complementary DNA (RNA extracts after reverse-transcription)
| Group | Species | Country | Collection Location / District / Region | N | Status | Body part | NA type |
| --- | --- | --- | --- | --- | --- | --- | --- |
| AA | *Cx. tritaeniorhynchus* | Bangladesh | Paba, Rajshahi | 10 | NBFF | Whole | gDNA |
| AB | *Cx. tritaeniorhynchus* | Bangladesh | Bagmara, Rajshahi | 7 | NBFF | Whole | gDNA |
| B | *Cx. tritaeniorhynchus* | Albania | Fier (Sop) | 16 | NBFF | Whole | gDNA |
| C | *Cx. pipiens* | Albania | Fier (Sop) | 16 | NBFF | Whole | gDNA |
| D | *Oc. caspius* | Albania | Fier (Sop) | 16 | NBFF | Whole | gDNA |
| E | *Cx. tritaeniorhynchus* | Albania | Fier (Sop) | 15 | NBFF | Abdomen | cDNA |
| F | *Cx. tritaeniorhynchus* | Albania | Fier (Sop) | 12 | BFF | Abdomen | cDNA |
| G | *Cx. tritaeniorhynchus* | Albania | Vlore | 10 | NBFF | Abdomen | cDNA |
| H | *Cx. tritaeniorhynchus* | Madagascar | Brickaville | 10 | NBFF | Whole | cDNA |
| I | *Cx. tritaeniorhynchus* | Madagascar | Farafangana | 9 | NBFF | Whole | cDNA |
| J | *Cx. tritaeniorhynchus* | Madagascar | Ihosy | 10 | NBFF | Whole | cDNA |
| K | *Cx. tritaeniorhynchus* | Madagascar | Maevatanana | 9 | NBFF | Whole | cDNA |
| L | *Cx. tritaeniorhynchus* | Madagascar | Miandrivazo | 9 | NBFF | Whole | cDNA |
| M | *Cx. tritaeniorhynchus* | Madagascar | Toamasina | 10 | NBFF | Whole | cDNA |
| N | *Cx. tritaeniorhynchus* | Madagascar | Tsaramandroso | 12 | NBFF | Abdomen | cDNA |
| O | *Cx. tritaeniorhynchus* | Madagascar | Tsaramandroso | 15 | BFF | Abdomen | cDNA |
| P | *Cx. antennatus* | Madagascar | Tsaramandroso | 14 | NBFF | Abdomen | cDNA |

#### 2.6.2 Data cleaning, quality control and filtering

Microbiome bioinformatics analyses were carried out on demultiplexed reads using “Quantitative Insights Into Microbial Ecology” (QIIME)2 Core (q2cli) 2020.2 distribution (104). Demultiplexed reads were divided according to extract type of gDNA (*16S* of all microbiota present) or cDNA (actively expressed microbial *16S*) (along with their respective blank control samples) and analyzed separately in downstream analysis. Reads were imported into QIIME2 and the V3-V4 primer and Nextera adapter sequences were removed using the “q2-cutadapt” plugin (105). Quality plots were generated and visualized using the “q2-demux summarize” command to assess and select optimal quality filtering parameters including truncation length for any adaptor sequence removal. Quality filtering, denoising and chimera removal was carried out using the “Diversive Amplicon Denoising Algorithm” (DADA) “q2-dada2” plugin (106) (“denoise-paired” command, gDNA: “p-trunc-len-f 231, p-trunc-len-r 229”; cDNA: “p-trunc-len-f 233, p-trunc-len-r 229”) to group Amplicon Sequence Variants (ASVs) within the data. The feature-table artifacts generated were filtered to exclude features present within the blank controls (“q2-feature-table filter-samples”).

#### 2.6.3 Taxonomic identification of features

Taxonomic assignment of ASVs was carried out using the “q2-feature-classifier” plugin (107) (“classify-sklearn” command (108)) with a pre-trained SILVA classifier (Naive Bayes classifier was pre-trained on the *16S rRNA* SILVA SSU v138 99% reference database (109), with the V3-V4 primers). The taxonomy classifier generated was used to remove mitochondrial and chloroplast ASVs from each feature table (“q2-taxa filter-table” plugin) to remove host and background non-relevant features. The samples were then filtered further (“q2-feature-table filter-features”) to remove features with frequencies below 100, and to only include the relevant samples for each comparative analysis. The taxonomic assignments were visualized using “q2-taxa barplot” to show relative taxonomic abundance across all individual samples (Figure 8).

#### 2.6.4 Alpha and Beta diversity analysis

Within the qiime2 phylogeny plugin, the “q2-phylogeny align-to-tree-mafft-fasttree” command was used, incorporating representative-sequence artifacts from each of the gDNA and cDNA groups (rep-seqs output from DADA2) to produce rooted phylogenetic trees for diversity analysis. For each comparison set, using the respective filtered feature tables, alpha and beta diversity analysis was conducted through the qiime2 diversity plugin, using “qiime diversity core-metrics-phylogenetic” (110). The sampling depth was selected from visualizing feature tables (“q2-feature-table summarize”) for each comparison, generating alpha-rarefaction visualizations (“q2-diversity alpha-rarefaction”) with “-p-max-depth” just over the median frequency per sample from the feature-table, and then by balancing the number of features, with the number of samples from each group retained (111). The diversity core metrics results were then generated and visualized using the relevant alpha or beta “group-significance” commands (112,113). Pairwise PERMANOVA tests with 999 permutations were used for comparisons between groups for the variable of interest and a significance level of P value <0.01 was used as the threshold. The metrics consulted for alpha (within-group) diversity were the Shannon diversity Index, Faith’s phylogenetic diversity and the Evenness. For beta (between-group) diversity the metrics consulted were the Bray-Curtis, Unweighted-Unifrac (pairwise) and Weighted-Unifrac.

#### 2.6.5 Differential abundance testing – ANCOM

To test for the presence of any differentially abundant taxa within each sample comparison group the analysis of composition of microbiomes (ANCOM) method was used within the qiime2 composition plugin (114). The “q2-composition add-pseudocount” command was used, followed by “q2-composition ancom” with the relevant variable selected for each comparison, to investigate if any association may be apparent. Results were visualized in volcano plots, and assessed through the test statistic, W, to determine significance.

### 2.7 Species-specific detection of *Wolbachia* and Multi-Locus Strain Typing

Amplification of *Wolbachia*-specific gene sequences was attempted using a range of assays targeting different *Wolbachia* genes in real-time or end-point PCR format. The conserved *Wolbachia 16S rRNA* gene was targeted using primers W-Spec-16S-F: 5’-CATACCTATTCGAAGGGATA-3’ and W-Spec-16S-R: 5’-AGCTTCGAGTGAAACCAATTC-3’ (end-point format, 438bp) (115), in addition to a primer set designed for real-time PCR (target length: 102bp, forward: 5′- CATACCTATTCGAAGGGATAG-3′, and reverse: 5′-TTGCGGGACTTAACCCAACA-3′) (116). The *Wolbachia* multi-locus strain typing (MLST) scheme (117) was employed to characterize *Wolbachia* strains using the sequences of five conserved genes as molecular markers to genotype each strain. In brief, 450–500 base pair fragments of the *gatB*, *coxA*, *hcpA*, *ftsZ* and *fbpA Wolbachia* genes were targeted. Primer sets used were as follows: gatB_F1: 5’- GAKTTAAAYCGYGCAGGBGTT-3’, gatB_R1: 5’-TGGYAAYTCRGGYAAAGATGA-3’, coxA_F1: 5’- TTGGRGCRATYAACTTTATAG-3’, coxA_R1: 5’- CTAAAGACTTTKACRCCAGT-3’, hcpA_F1: 5’-GAAATARCAGTTGCTGCAAA-3’, hcpA_R1: 5’-GAAAGTYRAGCAAGYTCTG-3’, ftsZ_F1: 5’-ATYATGGARCATATAAARGATAG-3’, ftsZ_R1: 5’-TCRAGYAATGGATTRGATAT-3’, fbpA_F1: 5’-GCTGCTCCRCTTGGYWTGAT-3’ and fbpA_R1: 5’-CCRCCAGARAAAAYYACTATTC-3’ (117). In addition, an alternative primer set targeting a 271bp fragment of the *ftsZ* gene sequence in *Wolbachia* strains from Supergroups A and B was used on selected samples; ftsZqPCR Forward: 5′-GCATTGCAGAGCTTGGACTT-3′ and ftsZqPCR Reverse: 5′-TCTTCTCCTTCTGCCTCTCC-3′ (118). PCR reactions and Sanger sequencing of *Wolbachia* MLST PCR products were carried out as previously described (119). Sequencing analysis was carried out in MEGA11 (87), using the methodology as described in section 2.3.2, with consensus sequences used to perform nucleotide BLAST (NCBI) database queries, and for *Wolbachia* gene searches against the *Wolbachia* MLST database (*http://pubmlst.org/wolbachia*). Phylogenetic analysis of MLST gene locus sequences was performed following methodology as described in section 2.3.3.

## 3 Results

### 3.1 Genetic diversity of *Cx. tritaeniorhynchus* populations

Sequences from a total of 69 *Cx. tritaeniorhynchus* specimens, originating from seven countries, spread across four continents (Figure 1), were generated in this study. Analysis of these *CO1* gene sequences, alongside all available reference sequences, comprised: a) maximizing coverage and comparison of the partial *CO1* gene fragment (primers from (78)) and concomitant species sequences, 253 sequences and 686 nucleotides; b) maximizing the length for comparison of the full *CO1* gene, 20 sequences and 1538 nucleotides; and c) maximizing the number of comparative sequences, with sufficient *CO1* fragment overlap, 1007 sequences and 414 nucleotides.

#### 3.1.1 Phylogenetic analysis and visualization

Phylogenetic analysis visualized the mosquito genetic variation across different populations and demonstrated the numerous clades of *Cx. tritaeniorhynchus* sequences, with geographic clustering to a certain degree (Figures 2 and 3). The phylogenetic trees from alignments a) and b) (Figure 2) show distinct grouping. The partial *CO1* phylogeny (Figure 2a) indicates four monophyletic groups for *Cx. tritaeniorhynchus*: Asia only; Asia, the Middle East and Eurasia; Africa and Europe; and Australia; with the concomitant species grouping separately. The detailed sub-tree of the Asia, Middle East and Eurasia group shows a distinct subclade, with sequences from the Middle East (Kuwait) and Eurasia (Georgia) diverging from the sequences from Asia within this group. The clade of African (Madagascar and Ghana) and European (Albania and Greece) sequences also includes two sequences from Bangladesh, but all other sequences from Asia group within one of the two main Asian clades. The most closely related sequences to *Cx. tritaeniorhynchus* are *Cx. sitiens*. Interestingly, two GenBank sequences recorded as *Cx. tritaeniorhynchus* (KM350638.1 and KM350640.1) are situated outside of the main *Cx. tritaeniorhynchus* phylogeny, and closer to the *Cx. sitiens* sequences. One of the sequences generated in this study from Tsaramandroso, Madagascar (MAD-16-TSA-CX-B1) could not be confirmed to be *Cx. tritaeniorhynchus* due to divergence to 94.58% identity when a BLAST search was performed, but interestingly the sequence most closely matched KM350640.1 in this search, and other sequences producing significant alignments are not *Cx. tritaeniorhynchus*, the next closest match being *Cx. dolosus*. These BLAST results and the phylogenetic tree placement suggests this specimen (and possibly KM350638.1 and KM350640.1) may sit outside of the *Cx. tritaeniorhynchus* species, but its ultimate discrimination has not been possible from the reference sequences currently available at this position of *CO1*. The full length *CO1* alignment (Figure 2b), although composed of fewer available sequences, indicates a similar geographic separation. Phylogenetic analysis of alignment c), maximizing the number of sequences included (Figure 3), demonstrates a more complex picture of the phylogenetic relationships between populations. The earliest common ancestors for this *Cx. tritaeniorhynchus* dataset were sequences from India and China, with divergence and a range of phylogroups then forming, with varying compositions of sequences from Asian countries-of-origin. These Asian-only clades include almost 70% (673/1007 sequences) of this dataset. The phylogeny then branches into several monophyletic groups containing sequences from other regions of the world. Sequences from Georgia obtained in this study, group most closely to sequences from Turkey and Kuwait, which in turn are most similar to a sequence from China, with this group branching from a sequence from Pakistan (Figure 3, blue inset sub-tree). Alongside further Asian clades, the next monophyletic group with more geographically diverse sequences include a group from Madagascar, branching from sequences from China, India and Pakistan (Figure 3, purple inset sub-tree). Another clade (Figure 3, green inset sub-tree), developing from a group of Indian sequences, includes the placements for sequences from Eurasia (Turkey), Africa (Ghana), Asia (Bangladesh) and Europe (Albania and Greece). The further divergence of this group then results in a group including sequences from Eurasia (Turkey), then branching to further sequences from Africa (Ghana and Madagascar), the Middle East (Saudi Arabia) and Europe (Albania and Greece). The sequence divergence suggested by the phylogenetic tree then continues to more Asian clades, a clade including sequences from Asia which then result in Eurasian (Turkey) and Middle Eastern (UAE) sequences, and finally branching to the placement of a clade containing further Asian sequences and the Australian sequences (Figure 3, yellow inset sub-tree). This group is most closely related to some of the available sequences from South Asia (India and Pakistan), with the sequence from Timor Leste (Southeast Asia, neighbouring Australia) also situated within the group of sequences from Australia. This phylogenetic analysis indicates that the lineages with the greatest extent of genetic distance – from the *Cx. sitiens* sequences included as an outgroup, and the suggested Asian *Cx. tritaeniorhynchus* ancestral sequences from India and China – appear to be those from Australia, followed by sequences from Madagascar and Europe (i.e. these sequences were placed furthest to the right of the phylogenetic tree).

**Fig 2:**
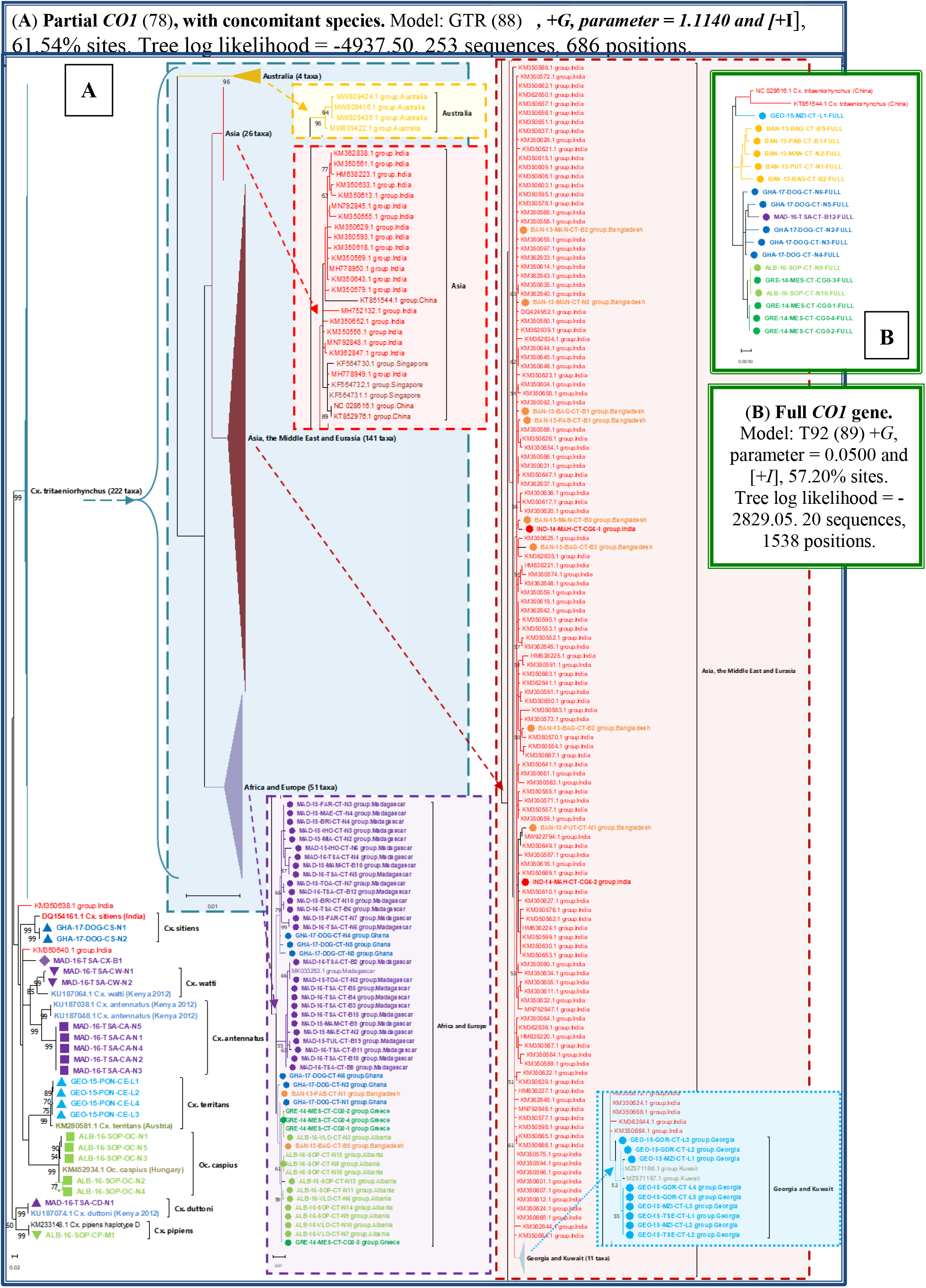
*Cx. tritaeniorhynchus* partial and full *CO1* maximum likelihood phylogenetic trees.

**Fig 3:**
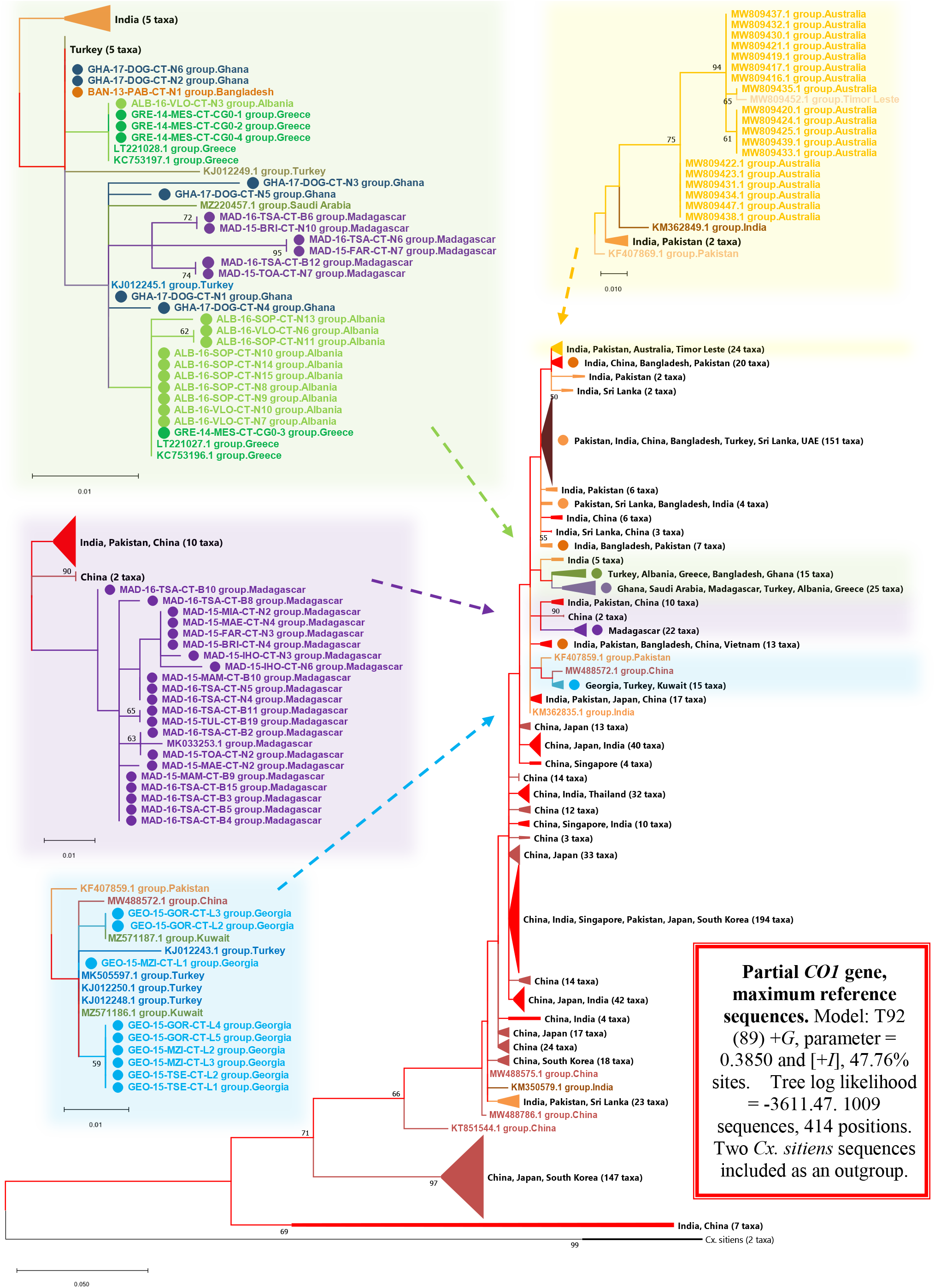
*Cx. tritaeniorhynchus CO1* phylogenetic tree with maximum reference sequences.

#### 3.1.2 Global genetic diversity metrics

The third alignment, c), maximizing the number of comparative sequences with sufficient *CO1* fragment overlap, was used to generate genetic diversity metrics (Table 2). Sequences were grouped according to country-of-origin and region, with the analysis including 21 countries and 6 regions. When all sequences were compared individually, across the 414 nucleotide positions in 1007 sequences, 139 variable sites (S) and 444 haplotypes (h) were identified. The overall haplotype diversity (Hd) was 0.97864, the average number of nucleotide differences (K) was 9.16425, and the nucleotide diversity per site (Pi) was 0.02214. The highest within-country nucleotide diversity per site was seen in South Korea (Pi = 0.03543) and lowest was in Greece (Pi = 0.00129). On a regional basis, the sequences from Asia produced the highest nucleotide diversity per site (Pi = 0.02203), and the sequences from Europe demonstrated the lowest within-region diversity (Pi = 0.00184).

#### 3.1.3 Haplotype networks and geographic haplotype mapping

*CO1* haplotype networks were constructed and visualized using alignments b) and c) (Figure 4). The full length *CO1* gene sequences (alignment b; 20 sequences, 1500 positions) (Figure 4a) produced a haplotype network suggesting a reasonably linear pattern of haplogroups, according to the sequence’s country-of-origin, with sequences from Asia (China and Bangladesh) clustering separately to sequences from Eurasia (Georgia), Europe (Greece and Albania), and Africa (Ghana and Madagascar). Haplogroups from Asia and Africa appeared the most divergent from one another, positioned at either side of the network.

**Fig 4:**
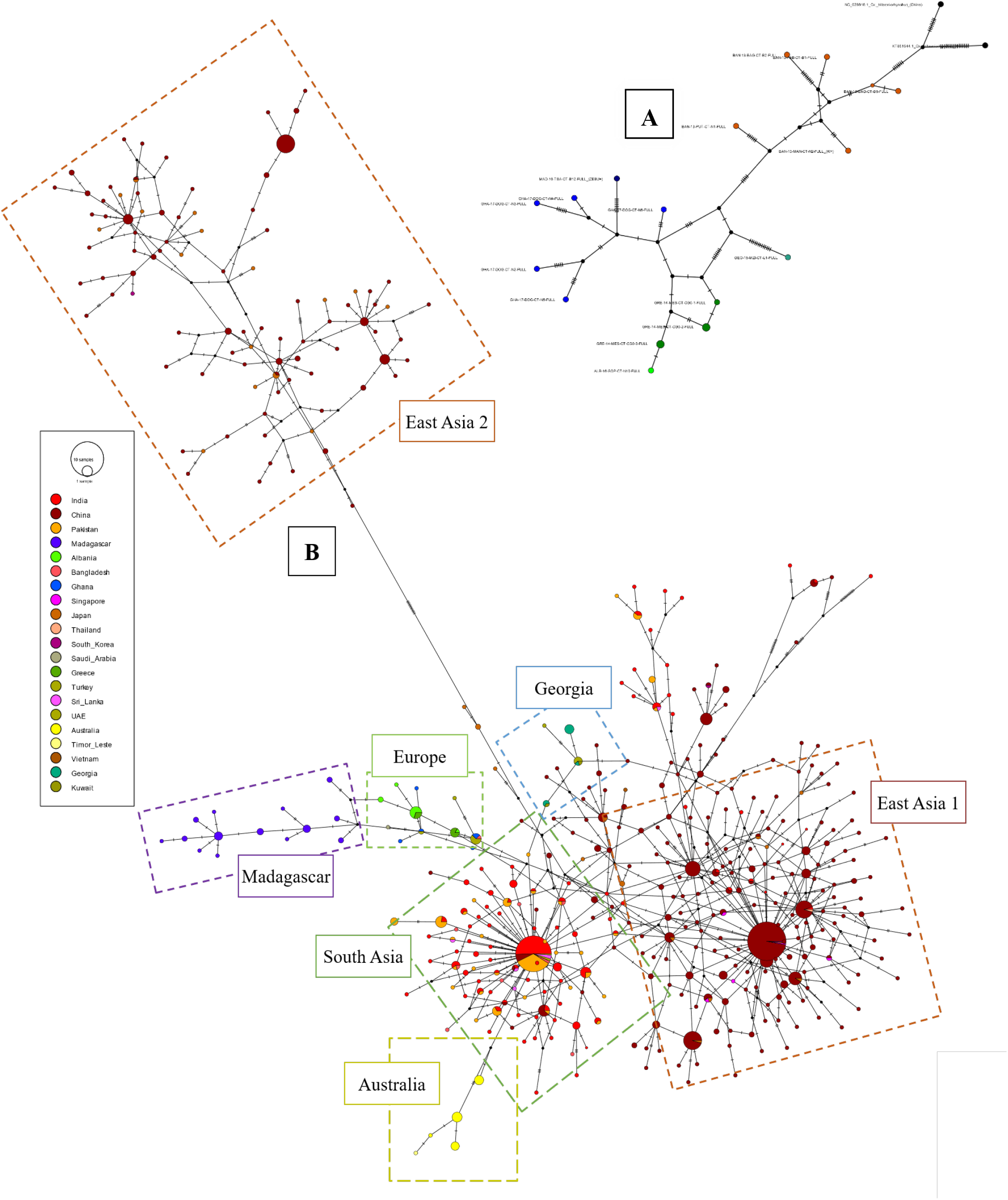
*CO1* haplotype networks for *Cx. tritaeniorhynchus*. (**A**) Full *CO1* gene haplotype network for *Cx. tritaeniorhynchus* (maximizing the length of sequences). (**B**) Global partial *CO1* haplotype network for *Cx. tritaeniorhynchus* (maximizing number of reference sequences). Haplotype networks were constructed using the TCS network method in PopArt (91) with nodes coloured according to country-of-origin.

The partial *CO1* gene sequences (alignment c; 1007 sequences, 414 positions) (Figure 4b) produced a more complex haplotype network, but where clear geographic clustering could still be seen between countries and regions. This haplotype network highlighted three major haplotypic groups from Asia, one mainly comprised of haplotypes from countries in South Asia, such as India, Pakistan and Bangladesh, and the other two comprising mainly haplotypes found in East Asia, such as from China, Japan and South Korea. One of these East Asian haplogroups diverges significantly from all other haplotypes. Haplotypes found in Australia and Timor Leste appear to be branching from the large South Asian cluster. The haplotypes from Georgia appear to branch from haplotypes present in Eurasia (Turkey) and the Middle East (Kuwait), sitting between the two large South, and East Asian foundational haplogroups. In a separate cluster, haplotypes from Europe (Greece and Albania) branch off from the main South Asian haplogroup, linked alongside some haplotypes present in Eurasia (Turkey) and Africa (Ghana). The haplotype present in Saudi Arabia also originates from this Eurasia/Africa branch, and the haplotypes present in Madagascar then extend and diverge further from the end of this branch.

The global *CO1* haplotype map (Figure 5), constructed through the analysis of alignment c, demonstrates the diversity and proportion of each of the 444 haplotypes which have resulted from the sequences available from each country. This visualization highlights the diversity of the haplotypes in each population and the geographic dispersal of haplotypes, in addition to the similarity in haplotypes between certain countries and regions.

**Fig 5:**
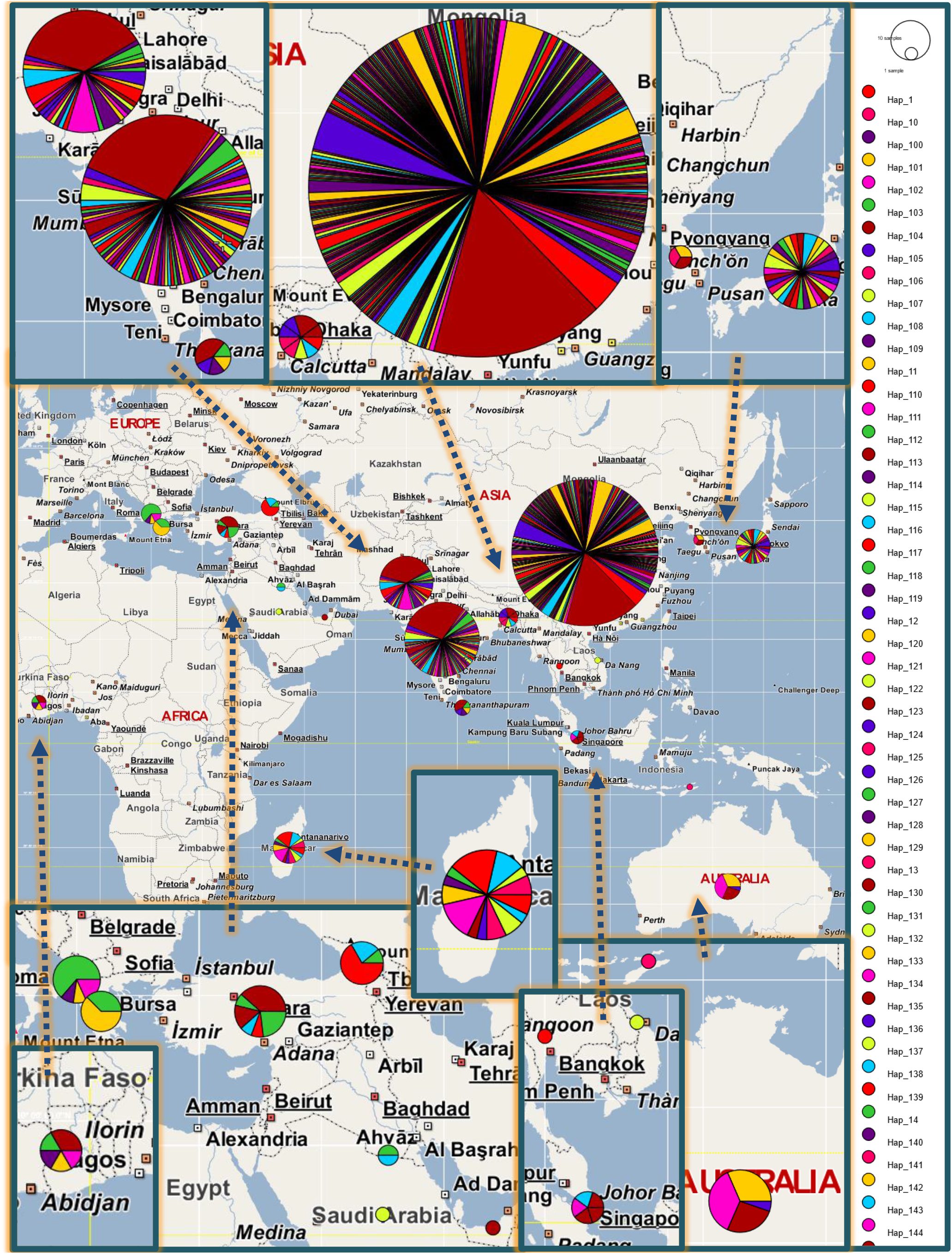
Global *Cx. tritaeniorhynchus CO1* haplotype map. PopART map view with *Cx tritaeniorhynchus CO1* Ref-seqs-Max data haplotype groups (geographic populations as traits added in nexus format)

#### 3.1.4 Pairwise comparison analysis for country-of-origin and region

Pairwise comparison analysis carried out on alignment c, enabled heatmaps to be generated for visualization (Figure 6). Analysis and visualization of the pairwise differences within and between populations of *Cx. tritaeniorhynchus* highlights the differences in genetic diversity between the different groups of sequences. The haplotype distance matrix highlights the divergence of some of the haplotypes, having a larger number of pairwise differences than the majority of the other haplotypes. This would seem to agree with the significant branching exhibited in the haplotype network. The average pairwise distances within and between groups, both at the country and regional level, highlights the sequences from Australia as having the greatest difference to other countries and regions, and the sequences from Asia, particularly South Korea, Japan and China, as having the greatest intra-group differences. The matrix of pairwise fixation index (*F*_ST_) indicates a fairly high genetic differentiation between populations in different countries and regions, particularly for the Australian, as well as the African and European groups. The divergence time between populations is also relatively lower for these populations, and highest for sequences from India.

**Fig 6:**
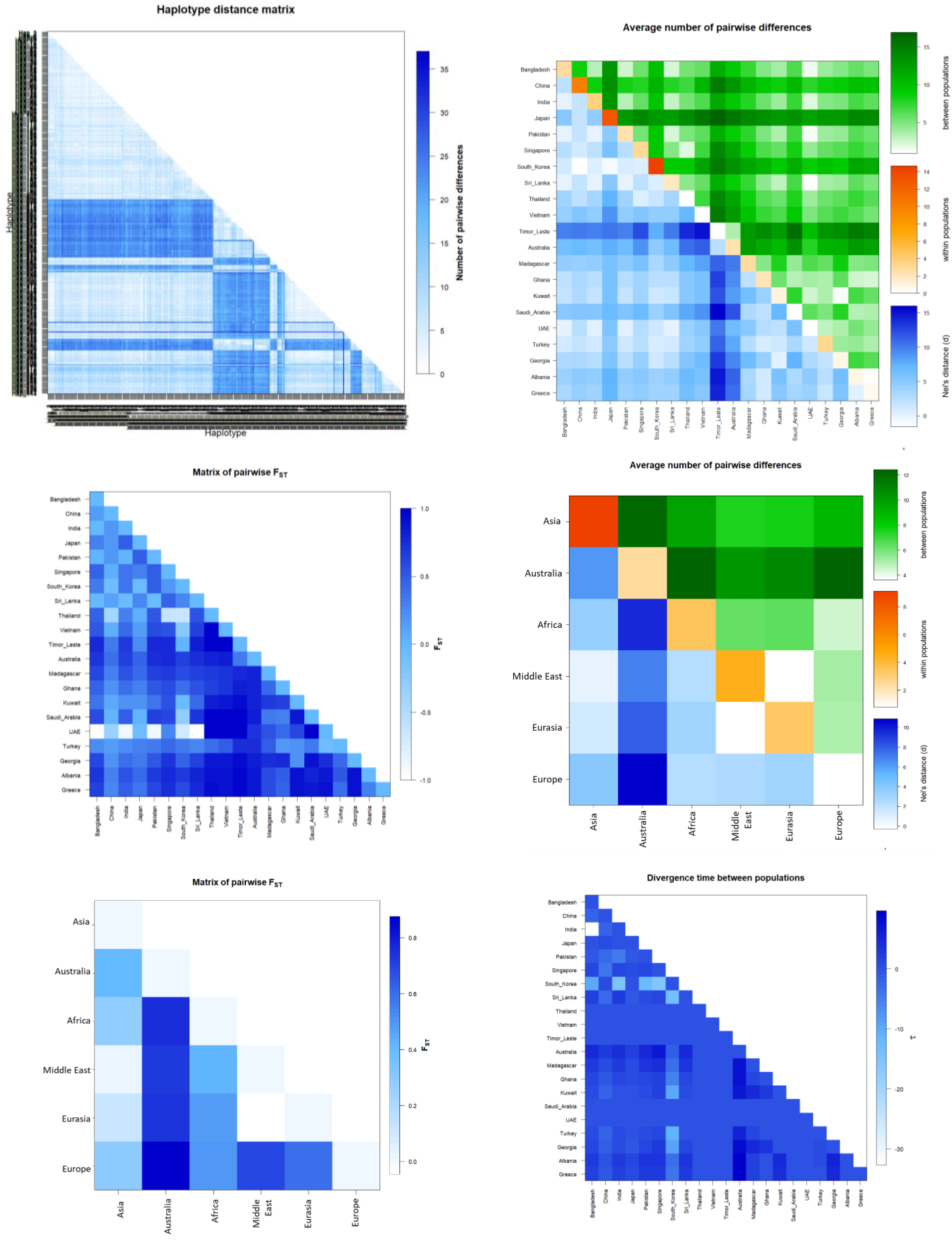
Global genetic diversity Country and Regional populations pairwise comparison heatmaps. (*CO1* Ref Seqs Max alignment, R visualizations from Arlequin analysis) (*F*_ST_ – pairwise fixation index: 0=two 1423 populations genetically identical, 1=two populations are genetically different, maximum genetic diversity between two 1424 populations)

### 3.2 JEV vector competence of colonized *Cx. tritaeniorhynchus* from Europe

A European colony of *Cx. tritaeniorhynchus*, established from wild larvae collected from Messolonghi in Western Greece, was fed an infectious blood-meal containing JEV (strain CNS138-11). After incubation for 14 days, high levels of JEV were detected in both the abdomen and head/thorax in all (28/28) surviving females, indicating the virus was successfully acquired and disseminated within these mosquitoes (Fig. 7). The mean qPCR Ct value for abdomen and head/thorax was 23.58 and 24.19 respectively (supplementary Fig S4). Our results also indicate a high level of JEV in the saliva collected from these individuals with 25/28 (89%) saliva extracts having detectable virus, with a mean qPCR Ct value of 26.93. This demonstrates that a high proportion of the mosquitoes were permissive to infections culminating in the excretion of viral material in saliva during feeding, as a proxy for the potential for onward transmission. A lower number of saliva samples had detectable virus after the 14-day incubation, compared to mosquito body parts, which might be expected, as the excretion of virus in the saliva is the final process in the infection pathway, following after viral acquisition and dissemination. However, this preliminary vector competence data clearly demonstrates this line of *Cx. tritaeniorhynchus* mosquitoes, colonized from wild-caught individuals collected in Greece, were highly competent vectors of JEV under these experimental conditions.

**Fig 7:**
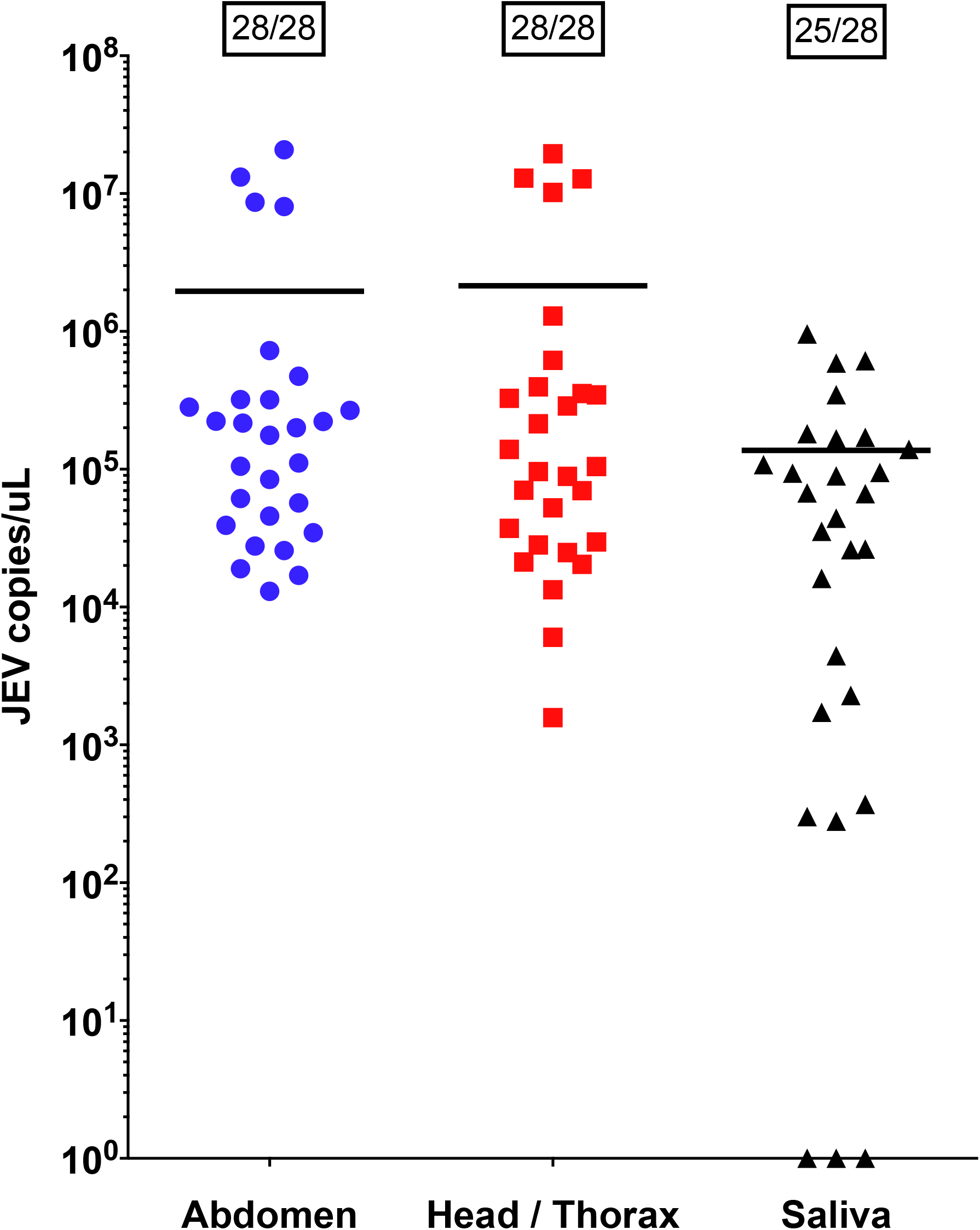
JEV vector competence experiment on European *Cx. tritaeniorhynchus* colonized from Greece. Scatter dot plot of quantitative PCR data. Horizontal bars represent mean JEV copies/μl per group. 1430 Boxed numbers show the number of JEV positive samples / total number of samples tested per 1431 group.

### 3.3 Virus detection and characterization

A subsample of field-collected specimens (which had been stored and preserved for RNA extraction, avoiding degradation of any potential viral RNA present,) were screened. An assay which utilizes degenerate primers targeting the flavivirus *NS5* gene (RNA-dependent RNA polymerase) and detects a range of both pathogenic and invertebrate-only flaviviruses, produced some positive results indicating flavivirus detection. Results from Greek adult specimens from field-collected *Cx. tritaeniorhynchus* larvae suggest the presence of a mosquito-only flavivirus. Sequencing analysis indicates this virus is likely to be a Cell Fusing Agent virus, within the non-pathogenic insect-only viruses group. Virus screening of *Cx. tritaeniorhynchus* specimens from Madagascar also identified the presence of RVFV as detailed previously (72).

### 3.4 Microbiome diversity

Bacterial *16S rRNA* gene amplicon sequencing was undertaken to determine the diversity of the mosquito microbiota, identify the bacteria of greatest relative abundance, the occurrence of bacterial species which have been implicated as potentially relevant to vectorial capacity and investigate variation in the composition and diversity of microbes present in *Cx. tritaeniorhynchus* between collection locations or countries, physiological states (blood-fed or non-blood-fed), or between *Cx. tritaeniorhynchus* and other concomitant mosquito species (Table 3, Figure 8).

**Fig 8.**
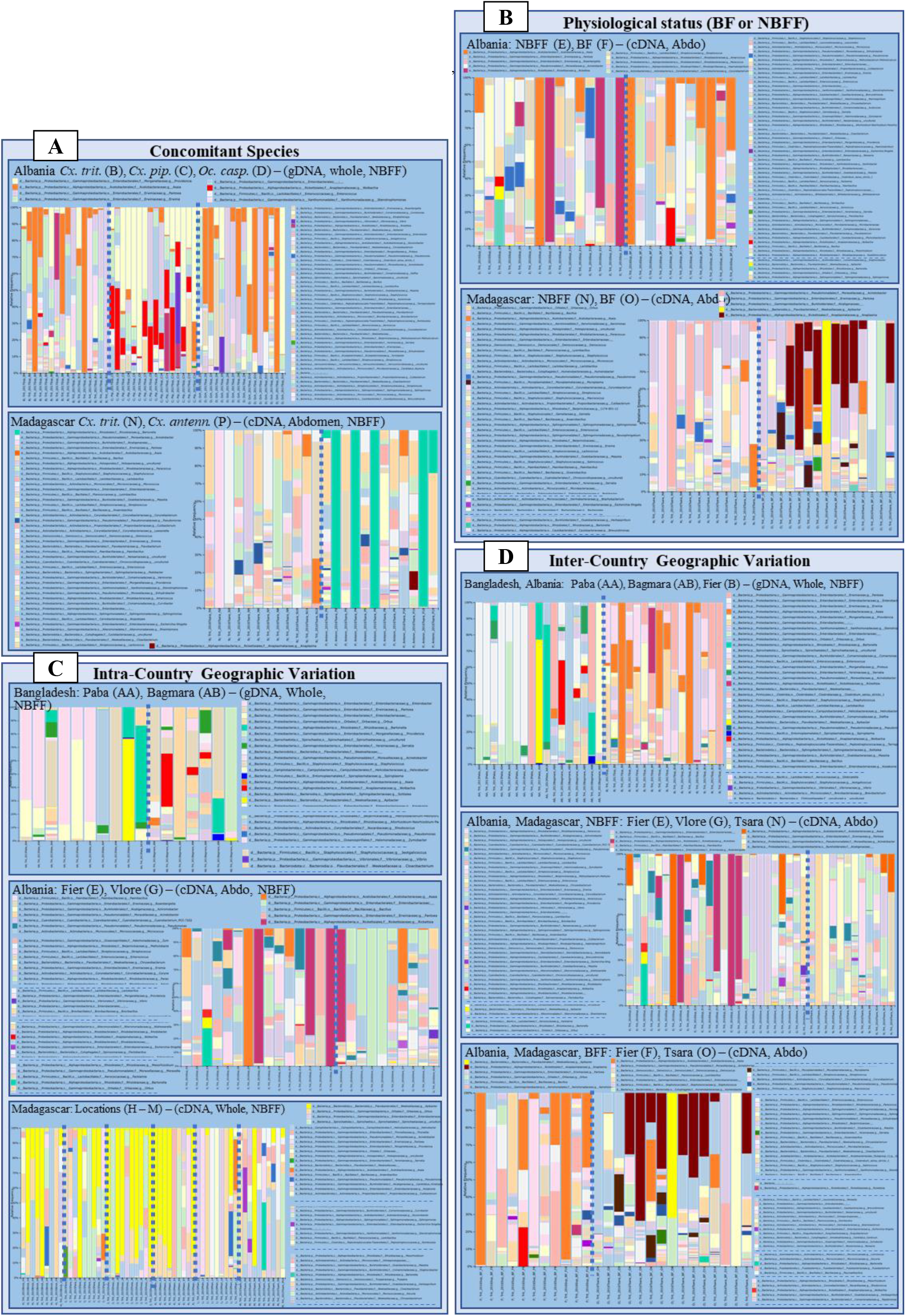
Microbiome analysis. Relative taxonomic abundance bar plots. *Wolbachia* in red, *Asaia* in orange, *Apibacter* in yellow, *Pseudomonas* in blue, *Serratia* in green, *Bartonella* in turquoise, *Vibrio* in dark purple, *Escherichia shigella* in light purple, *Rickettsia* in pink, *Anaplasma* in dark red and *Mycoplasma* in brown.

#### 3.4.1 The presence of *Wolbachia* and other microbes of relevance

Overall, taxonomic abundance analysis showed evidence for the presence of *Wolbachia* in some *Cx. tritaeniorhynchus* samples from Bangladesh, Albania and Madagascar. One sample from Bagmara, Bangladesh exhibited a relative abundance of *Wolbachia* comprising 39.77% of the total microbial composition, and two further specimens from the same location had 5.94% and 4.72% relative abundance respectively. A blood-fed specimen collected in Fier (Sop), Albania had a *Wolbachia* relative abundance of 22.50%. Three non-blood-fed samples from Albania, collected in Fier (Sop) and Vlore, also showed the presence of *Wolbachia* with relative abundances of 5.88%, 1.11% (Sop) and 1.32% (Vlore). These samples from Albania were cDNA samples, from RNA extractions of abdomens, rather than gDNA extracts from whole specimens as in Bangladesh. Concomitant Albanian *Cx. pipiens* mosquitoes (a species known to be naturally infected with the *w*Pip strain of *Wolbachia*) were shown to have variable relative abundances ranging from 0% (3 samples) to 38.24%.

In addition to *Wolbachia*, some other species of potential relevance to biocontrol were found, including *Asaia*, *Serratia*, *Pseudomonas* and *Apibacter* but presence and abundance levels were variable across different individuals and populations. For example, in cDNA whole-body non-blood-fed samples from Madagascar a substantial amount of *Apibacter* was found. Across the 57 specimens from six locations, there were 34 with some *Apibacter*, ranging from 0.05% to 99.87% relative abundance. In total, seven samples across this dataset (5 of the 6 locations) had *Pseudomonas*, ranging from 1.04% to 29.92%, one had *Serratia* present (20.38%) and three had Asaia, each of these from a different location (1.295% to 34.056%).

Of potential pathogenic importance, the presence of *Bartonella*, *Escherichia shigella*, *Vibrio cholerae*, *Anaplasma, Rickettsia, Mycoplasma, Enterobacter, Helicobacter* or *Providencia* was found in some individuals and populations, again of variable presence and abundance. For example, *Bartonella* was found in a sample from Fier (Sop), Albania (cDNA, non-blood-fed, abdomen) at a relative abundance of 27.45%. *Escherichia shigella* was found in two Albanian samples from Vlore (5.81% and 0.34% relative abundance), with this second sample also containing *Vibrio cholerae* at 6.28%. In non-blood-fed whole-body cDNA from Madagascar *Escherichia shigella* was identified in five specimens, across four locations, with abundance ranging from 0.41% to 30.00%. *Bartonella* was found in one specimen from Toamasina with 45.32% abundance and this specimen also had *Pseudomonas* at 8.63% abundance. In blood-fed *Cx. tritaeniorhynchus* from Tsaramandroso, Madagascar, *Anaplasma* was present in nine out of 15 blood-fed and none of the 12 non-blood-fed, with relative abundance ranging from 5.02% to 73.71%. Division down to taxonomic level 7 showed these ASVs were identified as *Anaplasma marginale*, *Anaplasma platys* and the rest classified within the *Anaplasma* genus. *Mycoplasma* was also identified in seven of the 15 blood-fed (0.33% - 21.61%) and none of the non-blood-fed. *Escherichia shigella* was found in four blood-fed (0.03% - 0.79%) and 2 non-blood-fed (0.84% - 1.91%).

#### 3.4.2 *Cx. tritaeniorhynchus* and concomitant species

For concomitant species comparisons, mosquitoes collected from Fier in Albania and Tsaramandroso in Madagascar were separately compared (Figure 8a). For Fier, Albania, gDNA samples from whole, non-blood-fed *Cx. tritaeniorhynchus* (n=16), *Cx. pipiens* (n=16) and *Oc. caspius* (n=16) specimens demonstrated variation in microbial composition. Alpha diversity within each species group showed no significant differences, however, beta diversity analysis showed clear differences between the species (Bray-Curtis p=0.001 and Weighted-Unifrac p=0.005). ANCOM identified *Wolbachia* as the only significant differentially abundant feature between the groups (W=265) due to the high abundance in *Cx. pipiens*. From Tsaramandroso, Madagascar, cDNA extracted from the abdomens of non-blood-fed female *Cx. tritaeniorhynchus* (n=12) and *Cx. antennatus* (n=14) demonstrated no significant difference in alpha or beta diversity and no significant differentially abundant taxa in ANCOM analysis.

#### 3.4.3 Comparing blood-fed and non-blood-fed female *Cx. tritaeniorhynchus*

To investigate the microbiota present in female *Cx. tritaeniorhynchus* of varying physiological states, comparisons were made of the microbiome results from cDNA extracted from the abdomens of non-blood-fed and blood-fed female specimens collected from Fier, Albania (non-blood-fed n=15, blood-fed n=12) and Tsaramandroso, Madagascar (non-blood-fed n=12, blood-fed n=15) (Figure 8b). For both countries, alpha- and beta-diversity and ANCOM highlighted no differences that were statistically significant.

#### 3.4.4 Variation between*Cx. tritaeniorhynchus* populations

In order to analyze the diversity of microbiota in *Cx. tritaeniorhynchus* from different populations, first the results from different locations within each country were compared to look at intra-country variation in Bangladesh, Albania and Madagascar separately (Figure 8c), and second, groups matched for other variables were compared between Bangladesh and Albania, and between Albania and Madagascar (Figure 8d).

For intra-country comparisons, in Bangladesh, specimens from two sites; Paba (n=10) and Bagmara (n=7), within the Rajashahi district (approximately 35km apart) were compared (gDNA from whole non-blood-fed females). No statistically significant differences were found through alpha- or beta-diversity, or ANCOM analysis, between the two locations. Within Albania, results from cDNA from non-blood-fed female abdomens from Fier (n=15) and Vlore (n=10) in south western Albania (approximately 35km apart) were compared. The differences between individuals within these groups were found to be significant (Faith’s phylogenetic diversity metric, p=0.0087), as well as between the groups (Unweighted-Unifrac, p=0.016) and *Enterobacteriaceae* was found to be significantly differentially abundant (ANCOM, W=156), with a higher abundance in Vlore than in Fier. For Madagascar, the microbiome data was generated from cDNA extracted from whole non-blood-fed females from 6 sites spread across Madagascar. These sites were Brickaville (n=10), Farafangana (n=9), Ihosy (n=10), Maevatanana (n=9), Miandrivazo (n=10) and Toamasina (n=10). Alpha diversity did not highlight any significant difference between individuals within the groups, but beta-diversity showed a difference between the locations (overall Weighted-Unifrac, p=0.003), with significance between Brickaville-Miandrivazo (p=0.012), Farafangana-Miandrivazo (p=0.012), Ihosy-Miandrivazo (p=0.001) and Maevatanana-Miandrivazo (p=0.004). ANCOM, however, found no significant differentially expressed taxa.

For inter-country comparisons, samples from Bangladesh and Albania (gDNA, whole-body, non-blood-fed) were compared and alpha-diversity demonstrated no significant difference between individuals within each group, whereas beta-diversity highlighted differences between each country, with Bray-Curtis (p=0.001) and Unweighted-Unifrac (p=0.001). ANCOM analysis showed there were several differentially abundant taxa between the groups from Bangladesh and Albania. The three taxa which were most statistically significant were two *Erwinia* species (W=235 and W=224) and *Asaia* (W=219), with their abundance in Albania much greater than in Bangladesh. For the comparison between Albania and Madagascar, one comparison was made between non-blood-fed and the other between blood-fed *Cx. tritaeniorhynchus* from the two countries (cDNA, abdomen samples). For non-blood fed, none of the alpha- or beta-diversity indexes showed significant results but ANCOM highlighted *Anaerobacillus* as a significant differentially abundant taxa (W=260), with higher abundance in Madagascar than Albania. This genus was present in nine of the 12 samples from Tsaramandroso, Madagascar (relative abundance range of <1% to 6.08%), present in four of 10 samples from Vlore, Albania, (<1% to 1.99%) and in none of the 15 samples from Fier (Sop), Albania. For the comparison between blood-fed females across the two countries, alpha-diversity showed no significant difference between the individuals in each group; from Fier (Sop), Albania (n=12) and Tsaramandroso, Madagascar (n=15). The Bray-Curtis beta-diversity metric showed a significant difference between the groups from each country (p=0.003) and ANCOM highlighted that reads classified in the *Bacillus* genus were significantly differentially abundant (W=266), with a relatively high abundance in samples from Madagascar, and absent from the blood-fed mosquitoes from Albania.

### 3.5 *Wolbachia*-specific detection and characterization

As *Wolbachia* was identified in samples using *16S rRNA* microbiome analysis, screening of further selected *Cx. tritaeniorhynchus* specimens using the *Wolbachia 16S* PCR (WSpec primers) revealed amplification of *Wolbachia* in certain samples from Albania, Greece, Madagascar and Bangladesh.

Confirmation of *Wolbachia 16S* amplification through Sanger sequencing was possible for some samples from Albania and Bangladesh (Table 4). Further analysis using the *Wolbachia* MLST gene loci showed a variable pattern of amplification and sequencing success, but partial MLST profiles could be obtained for *Wolbachia* positive samples from both Albania and Bangladesh (Table 5). The partial MLST allelic profile analysis, comparing the sequences obtained from *Cx. tritaeniorhynchus* specimens to those of the *Wolbachia* MLST database isolates exhibiting the closest or exact allelic matches at each locus, indicated these *Cx. tritaeniorhynchus Wolbachia* strains were different from one another, but were both placed within Supergroup B. Phylogenetic tree construction visualized these placements (Figure 9). Phylogenetic analysis of the *Wolbachia 16S* and the successful MLST gene loci sequences obtained, compared with reference sequences, confirms the strains from the individuals in Albania and Bangladesh are placed within Supergroup B. Although the strains do differ from one another where comparison was possible on the *Wolbachia fbpA* locus, they appear to be relatively closely related (Figure 9a).

**Table 4.**
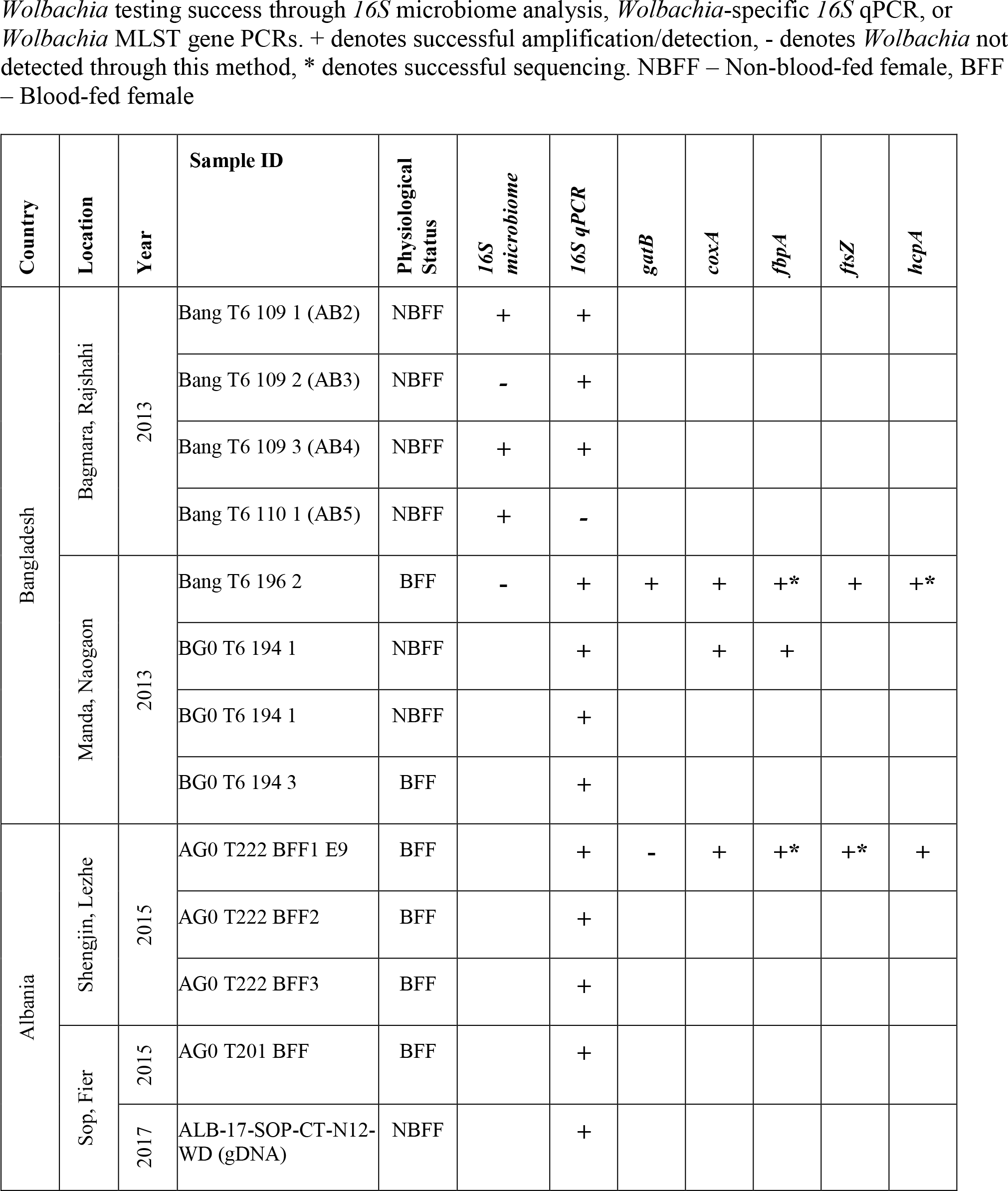
*Wolbachia* positive *Cx. tritaeniorhynchus* specimen details.

**Table 5.** *Wolbachia* partial MLST gene allelic profiles for resident strains in *Cx. tritaeniorhynchus* populations.

| Country and sample ID | *gatB* | *coxA* | *hcpA* | *ftsZ* | *fbpA* |
| --- | --- | --- | --- | --- | --- |
| Albania – AG0 T222 BFF1 E9 | NS | - | - | CM 280*<br>(0 diff)<br>SG B | EM 4<br>SG B |
| Bangladesh – BG0 T6 196 2 F9 | - | - | CM 12<br>(1 diff)<br>SG B | - | EM 27<br>SG B |

**Fig 9:**
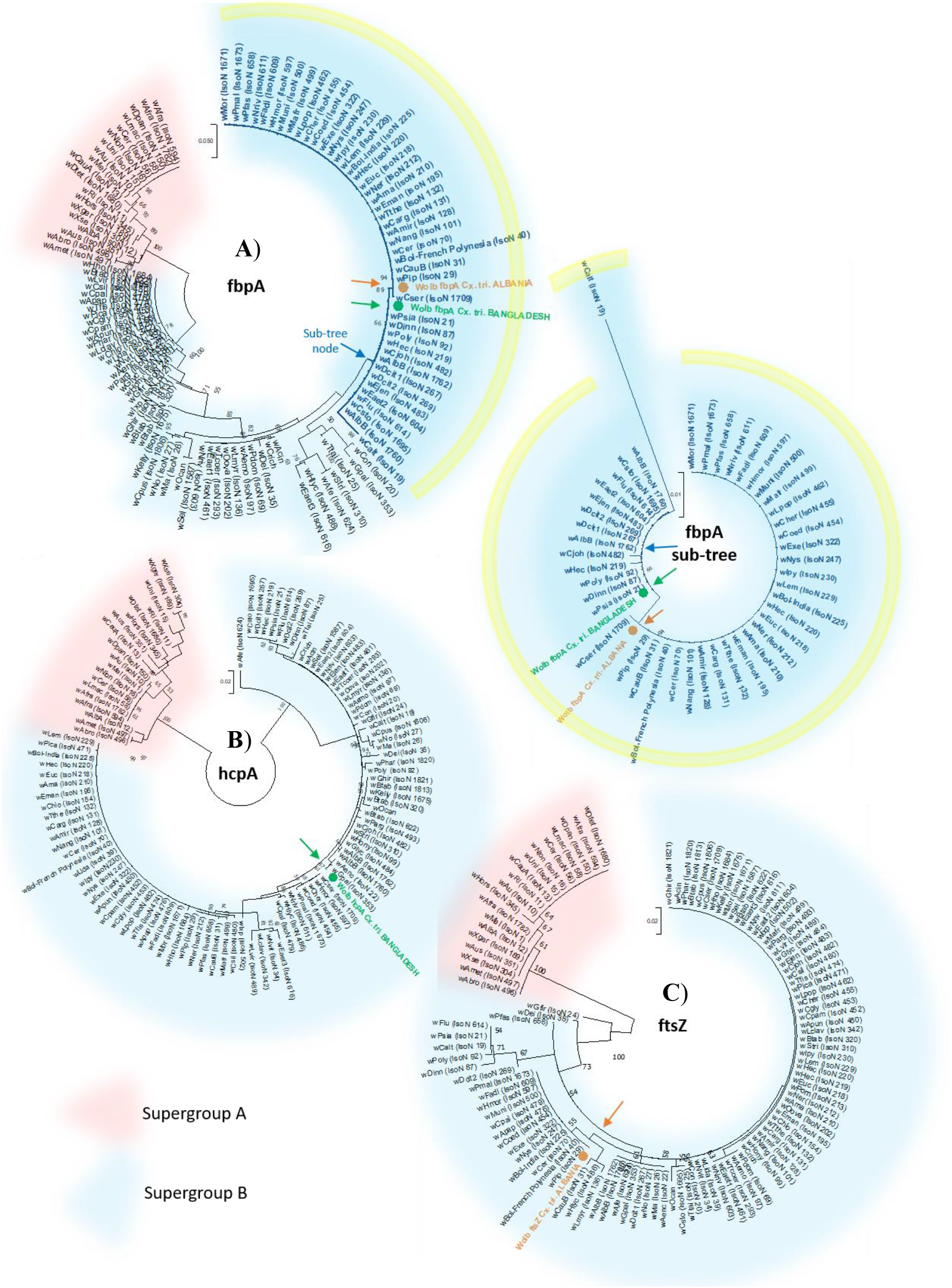
*Wolbachia* MLST gene phylogenetic trees. The T92 model (89) was used for all. *Wolbachia* Supergroups A and B are highlighted in red and blue respectively. The sequences obtained from *Cx. tritaeniorhynchus* from Albania and Bangladesh are shown in orange and green respectively. **A**)*fbpA* gene locus. Tree log likelihood = −1947.41 (+G, parameter = 0.2126). The analysis involved 113 nucleotide sequences and 429 positions. An *fbpA* sub-tree is also shown separately (yellow border) for more detailed visualization of the *Wolbachia fbpA* sequences obtained from *Cx. tritaeniorhynchus*. **B**) *hcpA* gene locus. Log likelihood = −1825.87. (+G, parameter = 0.3916). 111 nucleotide sequences and 444 positions. **C**) Partial *ftsZ* locus. Log likelihood = −535.33 (+G, parameter = 0.2600). 113 nucleotide sequences and 189 positions.

## 4 Discussion

There have been relatively few comparative studies on *Cx. tritaeniorhynchus* population diversity, or variation in vectorial capacity, with the majority characterizing *Cx. tritaeniorhynchus* populations within Asia (56,59,78,120–122). Historically this is where this species has had the greatest occurrence, abundance, and caused the greatest impact on human health as the major vector of JEV (14). However, the increasing occurrence of this species in other regions, and its potential role in pathogen transmission, as highlighted through its contribution as a major vector in the first incursion of RVFV outside of Africa (6), has warranted our comparative study of a broader range of *Cx. tritaeniorhynchus* populations.

Our analysis provides evidence for significant genetic diversity and adds to the existing debate of the correct classification (120,123,124) within the *Vishnui* subgroup and the *Cx. tritaeniorhynchus* species itself. The mitochondrial *CO1* gene has been the most frequently used for previous studies of *Cx. tritaeniorhynchus* (56,59,78,120,121,125) given it contains areas of highly conserved sequence, in combination with sufficiently diverse areas, allowing species discrimination and investigation of maternal inheritance patterns (125–127). The spatially diverse sequences generated in this study, alongside maximized comparisons with available reference sequences, has demonstrated the genetic differentiation occurring across the species current known geographic distribution. The lineages suggested by the phylogenetic analysis of the partial *CO1* gene with inclusion of the maximum number of reference sequences possible (alignment c), generating figure 3) is likely to be the most informative. Despite the need to compromise on the total coverage and number of sites includes in this alignment (due to the variable overlapping positions of the reference sequences available), this approach enabled the greatest quantity and geographic diversity of sequences to be compared. Our genetic diversity metrics quantified the genetic distances and divergence within population groups, identifying 444 haplotypes and 139 variable sites, with a haplotype diversity (Hd) of 0.97864 and a nucleotide diversity per site (Pi) of 0.02214. To our knowledge this is the first published study examining such a large and geographically diverse dataset – previous studies have identified 28 (125) and 303 (59) haplotypes, with the latter finding a Hd of 0.97 and Pi of 0.02434 which is comparable to our study. Analysis of regional population groups identified 412 haplotypes in Asia (n=909, Hd=0.97, Pi=0.02203), 4 in Australia (n=19, Hd=0.73, Pi=0.00565), 19 in Africa (n=34, Hd=0.96, Pi=0.00819), 4 in the Middle East (n=4, Hd=1.00, Pi=0.01087), 8 in Eurasia (n=22, Hd=0.86, Pi=0.00794) and 4 in Europe (n=19, Hd=0.64, Pi=0.00184). The only previous study for European *Cx. tritaeniorhynchus* found two haplotypes within the same population, collected in a single rice field in western Greece (70). Our samples from the same location identified two haplotypes which were also present in Albania, and a further two haplotypes were found in Albania only. The haplotype network and pairwise comparison analysis also indicated that (as expected) geographical location influences genetic diversity. For example, there was a distinct grouping of 14 haplotypes in Madagascar, none of which were found in any other countries or regions. The greater genetic distances of some groups, such as Australia, Madagascar, and Europe, is logical considering the extent of geographic distance and resulting genetic isolation of these populations from the original Asian lineages, but this also raises the question of what phenotypic effects genetic bottlenecks or selection pressures are having on this species during adaptation to new locations and environments.

This genetic data should enable a more accurate taxonomic classification of *Cx. tritaeniorhynchus*– particularly important as hybridization within species complexes (e.g. *Cx. pipiens*) can influence arbovirus transmission (128). Rapid, sensitive and specific molecular methods for the identification of *Cx. tritaeniorhynchus* are also paramount for vector control programs, and for surveillance to monitor this species in areas where it has historically been absent. There have been numerous recent reports of *Cx. tritaeniorhynchus* in countries where it had previously not been reported (2–5) highlighting a trend towards expansion of its known geographical range. Determining the phylogenetic origins of maternal lineages of *Cx. tritaeniorhynchus* can provide some insight into possible movement patterns when compared across countries and regions. During normal daily activity, *Cx. tritaeniorhynchus* are estimated to have an average flight distance of just under 70 meters, however, some studies have found that during long-distance wind-assisted dispersal, they are estimated to migrate between 200 and 500 kilometers (129). The adults overwinter and it is thought this species may use a combination of long-distance migration and hibernation *in situ*, as strategies to survive unfavorable conditions in temperate regions (130). The ability to disperse over such long distances and adapt to variable conditions is likely to provide more opportunities for range expansion and to increase gene exchange among different populations (59).

Despite the presence of *Cx. tritaeniorhynchus* first being reported in Europe – specifically Albania – in 1960 (131), further published European occurrence reports were scarce until the 2000s, with the species recorded in Greece; including from coastal marsh in Marathon near Athens (2), rice fields in Messolonghi, western Greece (70,132) and an urban area in Epirus, northwestern Greece (133). Recent extensive entomological surveys carried out in Albania have identified the presence of *Cx. tritaeniorhynchus* within multiple areas across the country. These reports highlight that this species appears to have become established in certain countries within southeastern Europe and may further expand its range in future. Entomological surveillance in Europe has also identified other invasive mosquito vector species such as *Aedes albopictus*, highlighting the risk of exotic vector species becoming established in the region (134). Concurrently, there has been an increasing trend of incursion, outbreaks and circulation of mosquito-borne arboviruses such as WNV in Europe, with many becoming established and endemic in multiple countries (135–137).

To our knowledge, this is the first study to assess JEV vector competence in a European population of *Cx. tritaeniorhynchus* and the results emphasize the possibility of future introductions and epidemics of vector-borne diseases. The previous detection of JEV RNA in mosquitoes and birds in Italy further reinforces this point (15,16). It was unfortunately not possible to generate comparative vector competence data between geographically dispersed populations as intended within the current study, due to the difficulties of obtaining, or colonizing, live mosquitoes from diverse populations in parallel. Most previous studies on *Cx. tritaeniorhynchus* JEV vector competence are difficult to directly compare to these results, due to differing infection and detection methods as they have developed over time. A study in the Republic of Korea resulted in 33-67% JEV transmission (via capillary tube saliva collection, or onward infection of chickens) (10) and in India, using ELISA in whole bodies, variable infection rates were reported, from 0-48% (60). The relatively lower infection rates seen in the Indian study may be, at least in part, as a result of reduced sensitivity of ELISA for virus detection or differences in the JEV infectious doses. With PCR-based JEV detection rates of 100% of abdomens (acquisition), 100% of head-thoraxes (dissemination) and 89% of salivary samples via capillary tube collections (transmission proxy), the current results, at minimum, provide no suggestion of the Greek *Cx. tritaeniorhynchus* being refractory to JEV under experimental conditions.

Although the direct extrapolation of laboratory vector competence experimental results to situations in the wild is likely to be imprecise given the complexity of transmission dynamics in wild populations, such data for local vector populations is an important component of vectorial capacity assessment. The species, genetics, age, fitness, immune response and microbiota of a mosquito will all influence its permissiveness to viral infection and speed with which it becomes infectious for onward transmission (42,43,138). Future studies performing vector competence experiments on adults directly from eggs or larvae collected from the field, and including time-course data to assess the extrinsic incubation period, would be logical next steps. Furthermore, environmental conditions such as temperature can affect transmission parameters within the vector, so future vector competence experiments could explore these factors, attempting to mimic more closely the current environmental niche of the populations. Establishing whether European populations of *Cx. tritaeniorhynchus* are competent vectors for other medically important arboviruses such as WNV and RVFV is also a priority. In addition, the assessment of host-vector associations, through blood-feeding patterns and host population densities, in differing localities would add valuable data for vectorial capacity assessment. From previous studies *Cx. tritaeniorhynchus* is generally thought to exhibit opportunistic host-seeking behaviours; primarily feeding on animals such as cattle and pigs but also feeding from a range of vertebrate hosts, including humans (139,140). A significant proportion of mixed blood-meals have also been found, an indication of feeding on multiple species of host within the same gonotrophic cycle (140). These feeding behaviours are likely to be context-dependent, but could greatly influence the transmission and infection dynamics, particularly in scenarios where *Cx. tritaeniorhynchus* could act as a bridge vector in zoonotic pathogen transmission networks (141).

The analyses of microbial diversity across *Cx. tritaeniorhynchus* populations in this study have also shown the variability that can occur. We found evidence for the presence of a non-pathogenic insect-only virus in some populations, suggesting there may be possibilities for co-infection to influence the dynamics of pathogenic viruses, and lead to potential effects on vectorial capacity and disease control strategies (41,52). The presence of insect-specific viruses in this species corresponds with other studies (62,64,66,142–144). The prevalence, transmission, evolution and impacts on pathogens of the insect-only viruses present in *Cx. tritaeniorhynchus* populations is an interesting avenue for further investigation and could prove particularly valuable if potential for utilization in biocontrol strategies could be explored and implemented.

Our microbiome analysis indicates that, as expected, there is a high degree of variability in the microbial composition both within and between populations of *Cx. tritaeniorhynchus*, as well as between other concomitant species. The taxonomic abundance analysis demonstrated the presence of *Wolbachia* in some *Cx. tritaeniorhynchus* individuals from different populations which, when present, showed variable levels of relative abundance from <1% up to approximately 40%. These results would indicate there is likely to be a variable, but mostly low level of *Wolbachia* infections present in populations from multiple countries and spread across several continents. The number of *Wolbachia* positive specimens from matching groups were not sufficient on this occasion to carry out analysis of the microbial composition and diversity between groups according to *Wolbachia* presence/absence.

The incidence and abundance of other microbes of potential relevance to biocontrol were also variable within and between populations. *Asaia* was present in all populations but not in all specimens and with highly variable relative abundance; dominating the microbiome of some individuals, while only contributing to a small proportion of the microbial composition when present in others. *Asaia* was differentially abundant in Albania when compared to Bangladesh (Figure 8d), but as *Asaia* can be environmentally acquired (145), it may depend on differing exposures in local habitats, rather than a country-wide distinction between populations. In the concomitant species comparison for Albania, between *Cx. tritaeniorhynchus*, *Cx. pipiens* and *Oc. caspius* (Figure 8a); *Cx. pipiens* with a greater abundance of *Wolbachia* (ANCOM, W=265), appeared to have a lower abundance of *Asaia* than the other two species sharing the same environment. A statistically significant difference wasn’t highlighted, and some individuals had both *Wolbachia* and *Asaia*, but seemingly at lower relative abundance than when either was identified without the other. Although speculative from this data, a reciprocal negative interference between the two has been found in previous studies (146,147). *Pseudomonas* and *Serratia* were also present in variable amounts across the different sample groups, however, they did not show a high abundance in any particular group, nor dominance of the microbiome in any individual.

The presence of *Apibacter*, with particularly high abundance in specimens from certain locations in Madagascar, some large contributions to relative abundance in these samples, and also present in a couple of individuals from Bangladesh and Albania, is interesting. *Apibacter* is a genus of bacteria classified within the Family *Weeksellaceae* which have been relatively recently first isolated and classified in 2016 from various bee species (148,149), as well as a strain being reported from house flies in 2019 (150) and one report in 2021 which found bacterial reads related to *Apibacter* in *Cx. fuscocephala* mosquitoes from Thailand (151). These bacteria are thought to be beneficial endosymbionts with characteristics of adaptation to the gut environment and a degree of host-specificity (152). They may also confer a degree of protection against pathogens, with a recent study finding an association between *Apibacter* in the microbiome of bees and decreased infection by a trypanosomatid gut parasite, *Crithidia bombi* (152,153). Although a far greater understanding would be required, this may suggest some parallels with *Wolbachia* which may be valuable to explore in future.

The bacterial genera *Anaplasma, Rickettsia, Bartonella, Vibrio, Helicobacter, Providencia, Mycoplasma* and *Escherichia* all contain some species and strains with pathogenic effects (154). For several, it was not possible to classify the ASVs beyond genus level and the species populating *Cx. tritaeniorhynchus* specimens may be non-pathogenic. For some, however, it was possible to identify to species, including *Escherichia shigella*, some strains of which can cause dysentery, and *Vibrio cholerae*, with certain strains causing severe cholera. These may have been present in the local aquatic environment and although surface sterilization prior to extraction and presence in cDNA from non-blood-fed abdomens would suggest active internal infections, it is possible these strains were environmentally acquired and attached to the chitin of mosquito exoskeletons. Their presence, pathogenic or not, does not imply the mosquitoes have capacity for onward transmission, although it may theoretically be possible for mosquitoes to mechanically disperse these bacteria from one local water source to another. *Anaplasma marginale* and *Anaplasma platys*, which can cause anaplasmosis in cattle and dogs respectively, are vector-borne pathogens, although they are mainly thought to be transmitted by ticks. A recent study of mosquito species in China, however, found a wide range of *Rickettsiales*, including *Anaplasma* spp., in mosquito species, including *Cx. tritaeniorhynchus*, from China (67). Phylogenetic analysis suggested a potential role for mosquitoes in vector-borne transmission of *Anaplasma marginale*, with other *Anaplasma* species suggested to be vertically transmitted, persisting as symbionts, with co-infections of differing *Anaplasma* species also occurring (67). Finding these bacteria at relatively high abundance in blood-fed mosquitoes in the current study, and not in the matched non-blood-fed mosquitoes may suggest these bacteria were present in the blood meals and not disseminated infections of the mosquitoes themselves. Even without vectorial capacity, however, mechanical transmission during blood-feeding may be possible. The high abundance in blood-fed-females would suggest a relatively frequent exposure to *Anaplasma*, particularly *Anaplasma marginale*, from their habits of feeding on cattle. If capable of mechanistic transmission, their opportunistic and multiple-host feeding behaviours may be a concern in this context. Bacteria from the *Bartonella* genus are also vector-borne, transmitted during blood-feeding, with ticks, fleas, lice and sandflies implicated as vectors in various transmission scenarios (155). Species from this genus can infect humans and animals, with bartonellosis causing a range of disease manifestations, depending on the infecting strain. *Bartonella* were identified in a few *Cx. tritaeniorhynchus* individuals from each country, – including gDNA and cDNA, non-blood-fed and blood-fed samples – and were highly abundant in some of these. Although in Madagascar, for the concomitant species comparison (cDNA, non-blood-fed, abdomens), *Bartonella* was found dominating the microbiome in 4 of the 13 concomitant *Cx. antennatus* specimens, but interestingly none was found in the matched group of *Cx. tritaeniorhynchus* from the same location.

In concordance with the microbiome results, *Wolbachia*-specific analysis and characterization also indicated the likely presence of low density *Wolbachia* strains in *Cx. tritaeniorhynchus*. This is evidenced through amplification produced from PCRs targeting the *Wolbachia* 16S rRNA gene and MLST gene loci, as well as sequencing data produced. Although amplification and sequencing success was variable, it was possible to obtain sequence data for some of the MLST genes and to carry out phylogenetic analyses to try to characterize the *Wolbachia* strains present in different populations further. Although some previous studies have not identified *Wolbachia* in this species (68,69,156,157), a study from Thailand (158) and recently from Singapore (159), reported *Wolbachia* in small numbers of *Cx. tritaeniorhynchus*. This mosquito is implicated as a vector of *Dirofilaria immitis*, which is one of the filarial nematode species with an obligatory symbiotic relationship with *Wolbachia*, requiring its presence for survival. However, the phylogenetic analysis carried out in this study clearly indicates the *Wolbachia* strains characterized here are not likely to be resulting from filarial infections, rather than from *Cx. tritaeniorhynchus* itself, due to the strain placements within Supergroup B (160). It remains to be determined whether these *Wolbachia* strains may influence reproductive success through the cytoplasmic incompatibility phenotype, whether they are vertically transmitted with high rates of maternal transmission, any impacts native *Wolbachia* strains may be having on *Cx. tritaeniorhynchus* population genetics and any interference with vectorial capacity. The apparently low infection rates and density of these strains may suggest they do not possess the beneficial characteristics most useful for biocontrol purposes and may not themselves demonstrate pathogen interference, as seen in some other studies on native strains (161). However, the presence of low-level natural *Wolbachia* strains in *Cx. tritaeniorhynchus* populations is unlikely to necessarily be prohibitive to the development of *Wolbachia*-based biocontrol strategies, such as through trans-infection of non-native strains with careful selection of advantageous strain characteristics, and it further demonstrates that *Wolbachia* is naturally present in some individuals from this species. Further analysis of larger sample numbers from diverse geographical areas is needed including non-PCR based methods such as microscopy to visualize bacteria in mosquito tissues (162). A greater understanding of the interactions of the other microbial constituents highlighted is also required to improve applicability of future biocontrol strategies.

## 5 Conclusions

This study provides the most comprehensive analysis of the genetic diversity of *Cx. tritaeniorhynchus* populations globally to date, including the largest number of geographically dispersed populations of *Cx. tritaeniorhynchus* so far. Despite the additional sequences generated in this study from spatially diverse samples, the remaining limitations of the relative quantities and uneven geographic distributions of the available *Cx. tritaeniorhynchus* sequencing data are still likely to be influencing the discriminatory power of the diversity analyses. Availability of a broader range of genetic data, with wide coverage of informative genes will be valuable in further understanding the phylogeography, divergence, range expansions and evolution of this species. The first full mitochondrial genome of *Cx. tritaeniorhynchus* has been published (58), and adding to the available genomic data in future with genomes obtained from geographically and genetically diverse *Cx. tritaeniorhynchus* populations will greatly expand the utility for comparison and potential for understanding this species and its contribution to vector-borne disease transmission. Until this study, to our knowledge, no arbovirus vector competence experiments have been carried out on European populations of *Cx. tritaeniorhynchus*. The results obtained here do nothing to suggest any reduction in vector competence as the species has expanded its range, adapted to new environments and genetically diverged from ancestral Asian lineages. The capability of populations from Europe or elsewhere to efficiently transmit JEV, or other arboviruses, is concerning and surveillance of this invasive species is needed. The microbial diversity results demonstrate evidence for the presence of likely low-density resident strains of *Wolbachia* in some diverse populations, as well as an insect-specific flavivirus associated with certain wild populations. Microbial community composition, and constituent relative abundances can be variable between individuals, locations, countries and continents, however, some similarities between different populations exist. The effects and interactions of endosymbiotic and potentially pathogenic microbes present in *Cx. tritaeniorhynchus* warrant further investigations and may augment the development of effective control strategies for this species. In addition to the current known possibilities for transmission of pathogens, future mosquito and pathogen range expansions with new geographic commonalities, and emergence of novel pathogens which *Cx. tritaeniorhynchus* is capable of transmitting, is a constant threat and one which may be further exacerbated in future with the effects of changing climate and other ecological parameters.

## 6 Conflict of Interest

The authors declare that the research was conducted in the absence of any commercial or financial relationships that could be construed as a potential conflict of interest.

## 7 Author contributions

C.L.J.: design, co-ordination, field-work, lab analysis, data analysis, manuscript writing, L.M.T.: Madagascar field-work, P.K.: Albania field-work, M.S.C.B.: Vector-competence experiments, I.L.: Greece field-work, J.O.: Ghana field-work, H.M.A.-A.: Bangladesh field-work, F.N.R.: Madagascar field-work, A.R.M.: Ghana field-work, M.S.A.: Bangladesh field-work, R.G.: Madagascar field-work, Y.A.A.: Ghana field-work, S.Bi.: Albania field-work, V.R.: Georgia field-work, S.Bo.: Madagascar field-work, M.B.: Vector-competence experiments, E.V.: Albania field-work, G.L.H.: microbiome sequencing and funding, T.W.: design, supervision, co-ordination, funding acquisition, manuscript revision. All authors assisted in drafting and approving the final manuscript.

## 8 Funding

C.L.J. and T.W. were supported by a Wellcome Trust/ Royal Society Sir Henry Dale Fellowship awarded to T.W. (101285/Z/13/Z).

P.K., S.Bi. and E.V. were supported by the Institute of Public Health, Albania and through the VectorNet framework, funded by the European Food Safety Authority (EFSA) and the European Centre for Disease Control (ECDC).

M.S.C.B. and M.B. were supported by grants from the BBSRC (BB/K018507/1), the MRC (ZK/16-041) and the BBSRC/ DEFRA (BB/W002906/1).

G.L.H. was supported by the BBSRC (BB/T001240/1 and V011278/1), a Royal Society Wolfson fellowship (RSWF\R1\180013), the UKRI (20197 and 85336), and the National Institute for Health Research (NIHR) (NIHR2000907).

M.S.C.B., M.B. and G.L.H. are affiliated with the NIHR Health Protection Research Unit (NIHR HPRU) in Emerging and Zoonotic Infections at the University of Liverpool in partnership with Public Health England (PHE), in collaboration with the Liverpool School of Tropical Medicine (LSTM) and the University of Oxford. M.S.C.B and M.B. are based at the University of Liverpool, G.L.H. is based at LSTM.

J.O. was supported through an MRC London Intercollegiate Doctoral Training Partnership Studentship.

V.R. was supported by a grant from the EFSA and the ECDC under the VectorNet framework (OC/EFSA/AHAW/2013/02-FWC1).

## 9 Acknowledgements

The authors would like to acknowledge and thank the following people for their assistance during this study: Eliot Hurn, Fara Raharimalala, Seth Irish, Laith Yakob, Elton Rogozi, Dusan Petric, and all the local people from each country where mosquitoes were collected.

## 13 Supplementary Material

### 13.1 RNA and DNA co-extraction method details

A method was tested and optimised to simultaneously, but separately, extract RNA and DNA from individual mosquito specimens in order to allow both RNA arbovirus screening and blood meal analysis of the DNA from the same individual. In brief, a *Cx. quinquefasciatus* colony from LSHTM was used to obtain specimens at various time intervals following a human blood-feed, to simulate the stages of blood meal digestion. Mechanical homogenisation using a Qiagen Tissue Lyser II (Hilden, Germany) with Qiagen 5mm stainless steel balls in Trizol (Invitrogen) was followed by the addition of chloroform to generate an aqueous upper phase containing RNA and a lower phase containing DNA and protein. The upper aqueous phase containing RNA was separated and ethanol was added, followed by continuation of the normal column-based RNA extraction procedure using Qiagen 96 RNeasy Kits (cat no. 74182) according to manufacturer’s instructions. RNA was eluted in 45 μl of RNase-free water and stored at −70°C. Proteinase K was added to the lower phase and DNA was extracted using Qiagen DNeasy Blood and Tissue kits according to manufacturer’s instructions. DNA extracts were eluted in a final volume of 100 μL and stored at −20°C. The method was carried out in either individual tubes, or 96-sample plate formats.

### 13.2 PCR and Sanger sequencing method additional details for molecular species identification and mosquito genetic diversity sequence generation

PCR products were separated and visualized using 2% E-Gel EX agarose gels (Invitrogen) with SYBR safe and an Invitrogen E-Gel iBase Real-Time Transilluminator.

PCR products were submitted to Source BioScience (Source BioScience Plc, Nottingham, UK) for PCR reaction clean-up, followed by Sanger sequencing to generate both forward and reverse reads. Primers used for sequencing were the same as used in the original PCR amplification for generation of products.

---

**Fig. S1.**
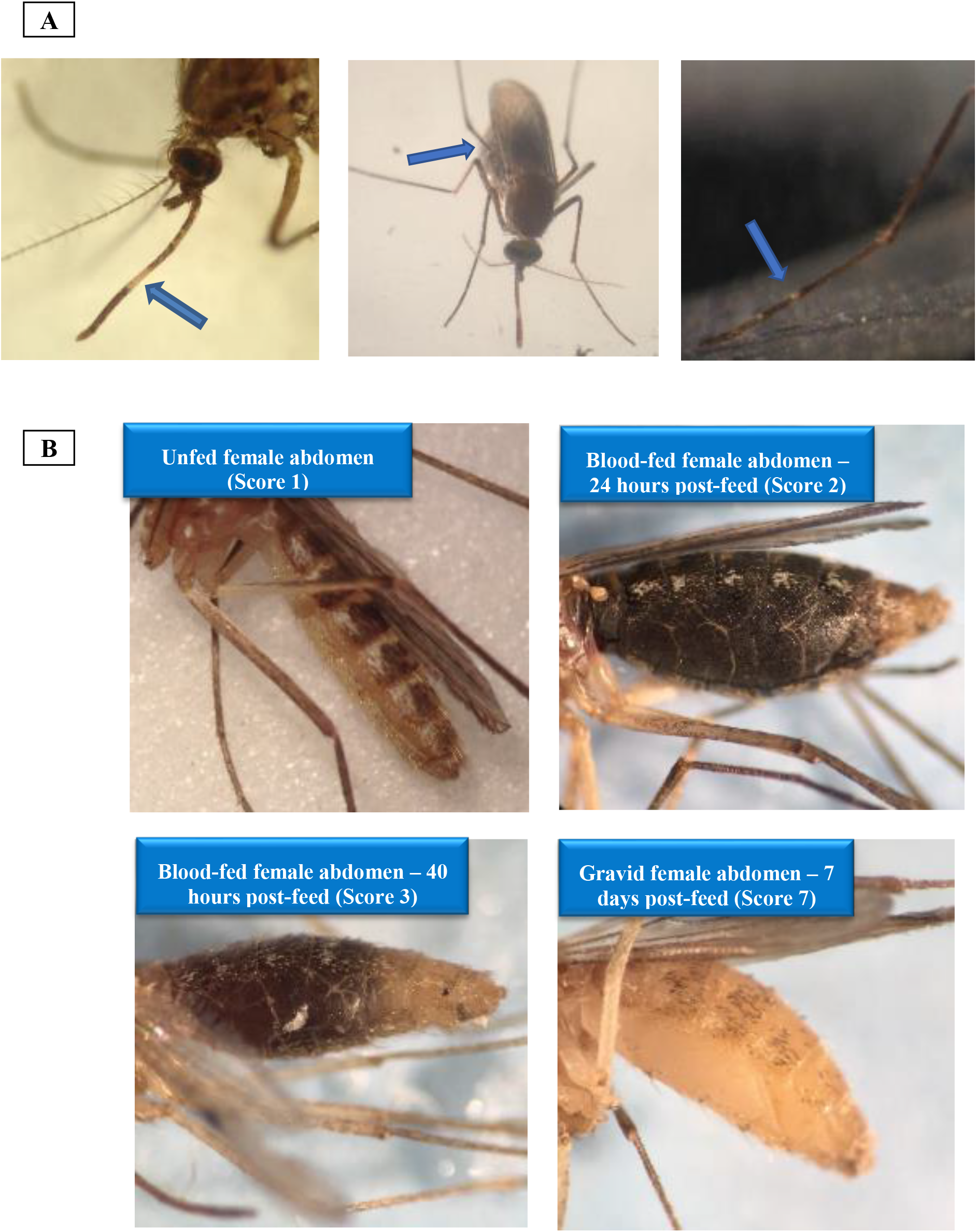
Morphological identification of *Cx. tritaeniorhynchus* and Sella score for blood-fed females.

**Fig. S2:**
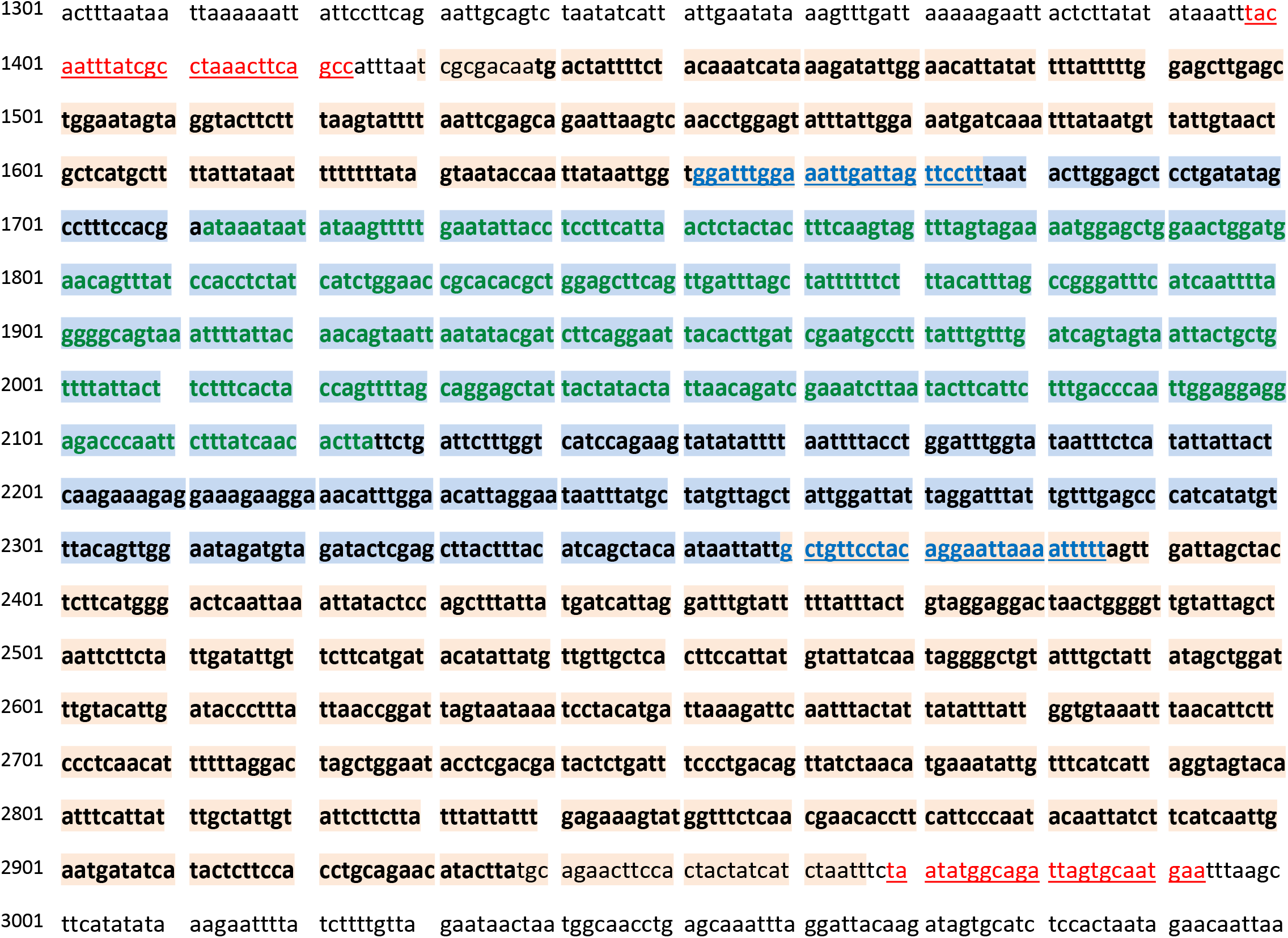
*CO1* primer sets and sequence alignment positions. Excerpt of nucleotide sequence from the complete mitochondrial reference genome of *Cx. tritaeniorhynchus* (NC_028616). Red, underlined text – primer binding sites of the full length *CO1* primer combination; Orange background – nucleotides comprising the full *CO1* gene; Blue, underlined – Kumar *et al*. (78) primer binding sites; Blue background – Position of alignment (a) of sequences spanning the length between the Kumar *et al*. primer set binding regions; Text in bold – Location of alignment (b) maximizing the length and with almost full *CO1* gene coverage; Green text – Location of alignment (c) maximizing the number of *Cx. tritaeniorhynchus CO1* sequences included.

**Fig S3:**
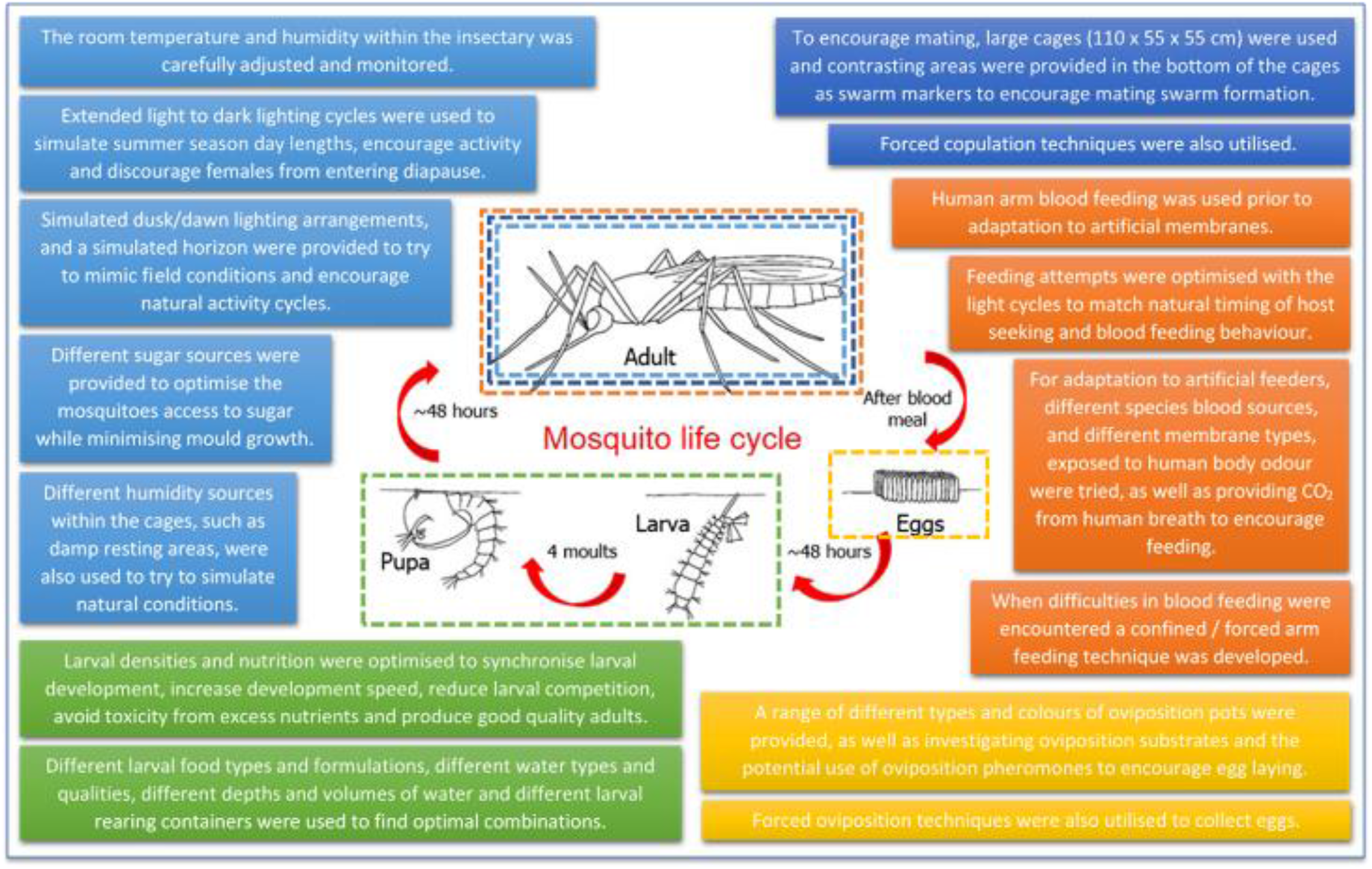
Optimisation of techniques for colonization of *Cx. tritaeniorhynchus* originating from Greece for vector competence experiment.

**Fig S4:**
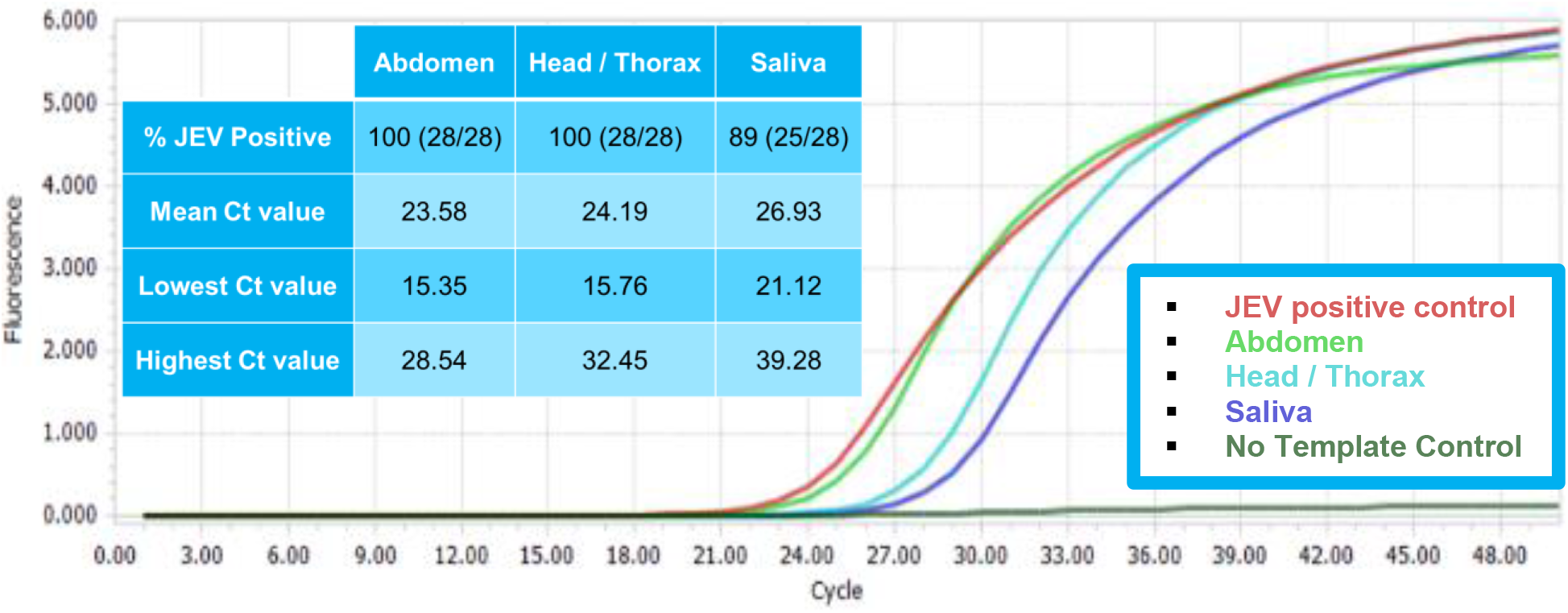
JEV vector competence qPCR summary. Representative amplification plots of samples from one individual, with controls, and summary qualitative results (table inset).

## Notes

### Competing Interest Statement

The authors have declared no competing interest.

